# CellEKT: A robust chemical proteomics workflow to profile cellular target engagement of kinase inhibitors

**DOI:** 10.1101/2024.10.01.616061

**Authors:** Joel Rüegger, Berend Gagestein, Antonius P. A. Janssen, Alexandra Valeanu, Alger Lazo Mori, Marielle van der Peet, Michael S. Boutkan, Bogdan I. Florea, Alex A. Henneman, Remo Hochstrasser, Haiyan Wang, Paul Westwood, Andreas Topp, Patricia M. Gomez Barila, Jan Paul Medema, Connie R. Jimenez, Bigna Woersdoerfer, Stephan Kirchner, Jitao David Zhang, Uwe Grether, Arne C. Rufer, Mario van der Stelt

**Affiliations:** Department of Molecular Physiology, Leiden Institute of Chemistry, Leiden University & Oncode Institute, Einsteinweg 55, 2333 CC Leiden, The Netherlands; Department Medical Oncology, OncoProteomics Laboratory, Cancer Center Amsterdam, Amsterdam UMC, Location VUmc, Amsterdam, The Netherlands; Roche Pharma Research & Early Development, Roche Innovation Center Basel, F. Hoffmann-La Roche Ltd., 4070 Basel, Switzerland; Bio-Organic Synthesis, Leiden Institute of Chemistry, Leiden University, Einsteinweg 55, 2333 CC Leiden, The Netherlands; Laboratory for Experimental Oncology and Radiobiology, Center for Experimental and Molecular Medicine, Cancer Center Amsterdam, Amsterdam UMC, University of Amsterdam and Oncode Institute, Meibergdreef 9, 1105 AZ Amsterdam, The Netherlands

**Keywords:** CellEKT, cellular target engagement, endogenous kinome profiling, XO44, IC_50_, drug selectivity, broadspectrum kinase probes, chemical proteomics, ABPP

## Abstract

The human genome encodes 518 protein kinases that are pivotal for drug discovery in various therapeutic areas such as cancer and autoimmune disorders. The majority of kinase inhibitors target the conserved ATP-binding pocket, making it difficult to develop selective inhibitors. To characterize and prioritize kinase-inhibiting drug candidates, efficient methods are desired to determine target engagement across the cellular kinome. In this study, we present CellEKT (Cellular Endogenous Kinase Targeting), an optimized and robust chemical proteomics platform for investigating cellular target engagement of endogenously expressed kinases using the sulfonyl fluoride-based probe XO44 and two new probes *ALX005* and *ALX011*. The optimized workflow enabled the determination of the kinome interaction landscape of covalent and non-covalent drugs across over 300 kinases, expressed as half maximum inhibitory concentration (IC50), which were validated using distinct platforms like phosphoproteomics and NanoBRET. With CellEKT, target engagement profiles were linked to their substrate space. CellEKT has the ability to decrypt drug actions and to guide the discovery and development of drugs.

**Graphical abstract:** 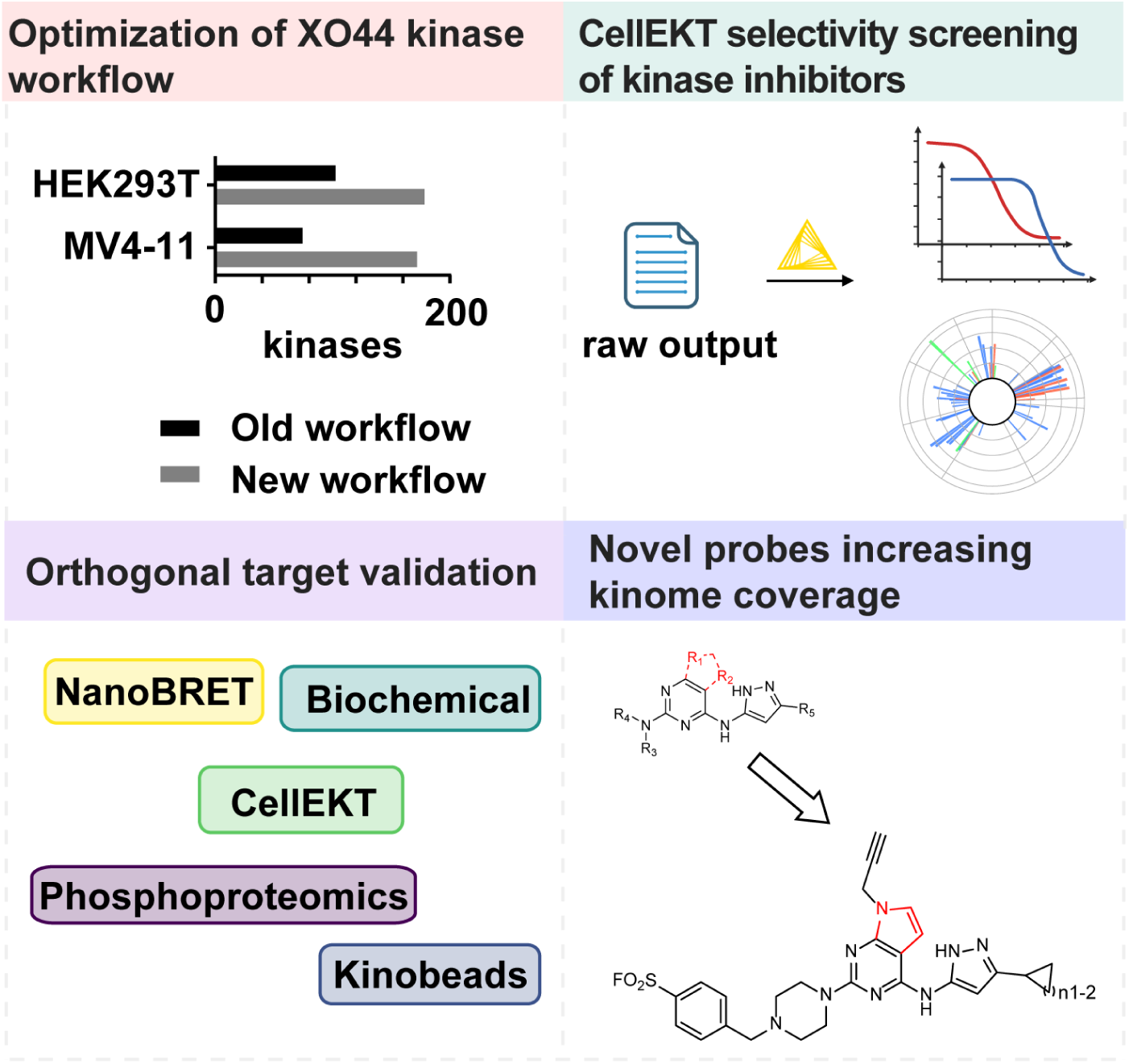

## Introduction

Kinases are important enzymes that phosphorylate proteins or cellular metabolites, regulating protein function, signaling pathways, and cellular function. The human genome encodes 637 kinases with 518 targeting proteins and others phosphorylating lipids, carbohydrates, or other metabolites.^1,2^ Kinase dysregulation is linked to diseases like cancer, inflammatory conditions and autoimmune disorders, making kinases vital drug targets.^3^ The discovery of kinase inhibitors is a major focus in both academic and industrial research, culminating in the 80 FDA-approved kinase inhibitors as of writing.^4^ Many inhibitors target the conserved ATP-binding site in kinases, often leading to off-target interactions resulting in adverse side-effects.^5^ Due to toxicity and efficacy issues, most kinase inhibitors fail in clinical trials. Only a 41% success rate from phase I to approval was noted.^6^ Thus, understanding kinase inhibitors’ interactions with their targets is crucial, especially as they are developed for non-oncological conditions.^3^

Binding and activity assays with purified kinases or their catalytic domains, along with mass spectrometry-based methods, employing irreversible ATP-biotin probes,^7^ photoaffinity-based probes^8^ and kinobeads,^9–11^ are commonly used to study the selectivity profile of kinase inhibitors. However, these assays fail to replicate the complex cellular environment, which can influence enzyme activity through post-translational modifications, cellular localization, substrate levels, and protein-protein interactions.^12,13^ Kinase inhibitors targeting the ATP binding site must also compete with cellular ATP, present at 2-8 mM in mammalian cells, which is two to three orders of magnitude higher than the enzyme K_M_, frequently used in biochemical assays under balanced conditions.^14,15^ This discrepancy can lead to different cellular target engagement profiles compared to those obtained from biochemical and lysate-based assays.

To address these limitations, several methods have been developed to study target engagement in living cells. The cellular thermal shift assay (CETSA)^16^ is a powerful technology for full proteome drug target screenings^17^ and has been successfully applied to *in vivo* studies and whole blood samples.^18^ However, low-abundant, membrane-bound or unstable targets are challenging to assess with CETSA, which may result in overlooked drug interactions.^16^

Another technology for cellular selectivity screening is the energy transfer (NanoBRET) assay, which has been applied across different enzyme classes.^19,20^ NanoBRET has advanced the selectivity screening of kinase inhibitors, assessing inhibitor specificity across a spectrum of 178 kinases in a 96-well format in transfected HEK293 cells.^15^ Refinements in the NanoBRET platform has allowed the substitution of six BRET tracers with a single one, and extended the coverage to 192 kinases^21^ with the capability to perform automated screenings in a 384-well format.^22^ Despite these advancements, this platform relies on overexpressed, transfected constructs, limiting its applicability across different biological systems such as various cell lines, organoids, animal models, or primary human cells.^23,24^

Chemical proteomics has emerged as a powerful technique to assess the target interaction landscape of small molecules using chemical probes in a native biological context. XO44 was the first kinase probe developed for this purpose.^25^ It is a broad-spectrum kinase inhibitor with a sulfonyl fluoride warhead that reacts covalently with a conserved lysine in the kinase pocket. It also includes a bioorthogonal ligation handle for attaching a fluorophore for visualization or a biotin for affinity enrichment. In the original work, XO44 was used to identify 133 kinases in Jurkat cells and to map the cellular target-interaction landscape of dasatinib. Additionally, XO44 was used to study kinome expression in lenvatinib-resistant hepatocellular carcinoma cells, identifying CDK6 as a resistance driver.^26^ Modifications to the scaffold of XO44 have included a salicylaldehyde warhead to determine the target profile of a CDK inhibitor in mice,^27^and substituted aryl fluorosulfates to profile the kinase lysine reactivity.^28^ These studies show the value of chemical proteomics in establishing target engagement studies in native biological settings. Although XO44 is commercially available, thereby facilitating broad usage of this probe, no best practices for its use have been reported.

Here we report an optimized chemical proteomics workflow called CellEKT (Cellular Endogenous Kinase Targeting) for profiling cellular target engagement of kinase inhibitors using XO44 and two new chemical probes. By employing CellEKT, we profiled the cellular interaction landscape of the FDA-approved kinase inhibitors dasatinib, midostaurin and three Bruton’s tyrosine kinase inhibitors. The targets of these drugs identified by CellEKT were validated through diverse methodologies, including biochemical assays, NanoBRET technology, and phosphoproteomic analysis. Finally, by leveraging resources of protein kinase substrate specificity ^29,30^ and computational tools^31^, we show that CellEKT helps to define substrate space and elucidate drug mechanisms and off-target effects. In summary, CellEKT is a new tool to gain a comprehensive understanding of therapeutic and toxicological mechanisms of actions for kinase drug discovery.

## Results

### Optimization of sample preparation

The general chemical proteomics workflow is composed of five steps (1-5) that present individual challenges, as shown in Figure 1A. To improve these steps, covalent labeling by the probe was visualized by ligation to a fluorophore azide. The separation of proteins was done using sodium dodecyl sulfate polyacrylamide gel electrophoresis (SDS-PAGE), followed by fluorescence scanning, to compare treatment conditions. The pull-down conditions were investigated by ligating probe-bound proteins to the trifunctional 5-carboxytetramethylrhodamine biotin-azide (TAMRA-biotin-azide), allowing for the tracking of labeled proteins in supernatant and pull-down bead fractions.^32^ In step 1 of the workflow the cell density was optimized by treating various concentrations of THP-1 cells with XO44 at a concentration of 1 µM to avoid the risk of probe depletion. Integration of the fluorescent signal on the SDS-PAGE gel showed that a cell density of 1×10^6^ cells/mL is ideal using 1 µM XO44 over 30 min of incubation (Figure S1A). In step 2, cell lysis was optimized by investigating the effect of post-lysis labeling. Initial tests showed that probe labeling continued after cell lysis if the lysate was not kept on ice (Figure S1B). The sudden drop in ATP concentration was suspected to contribute to the post-lysis labeling. Post-lysis labeling was eliminated by increasing SDS concentration to 1%. In step 3, the copper-catalyzed azide–alkyne cycloaddition (CuAAC) ligation was optimized using tris(2-carboxyethyl)phosphine (TCEP) instead of sodium ascorbate (Figure S2A and S2B). The increased SDS concentration reduced reactive oxygen species (ROS) related band smearing (and S2C).^33^ Finally, in step 4 the pull-down of biotinylated proteins was optimized (Figure S2D).^34,35^ Residual protein labeling in the supernatant was examined by adjusting both the incubation time and the quantity of agarose beads (Figure S3A-D). To further decrease bead usage without compromising signal detection, incorporating unmodified agarose beads helped to retain the proteins post-washing.

**Figure 1:**
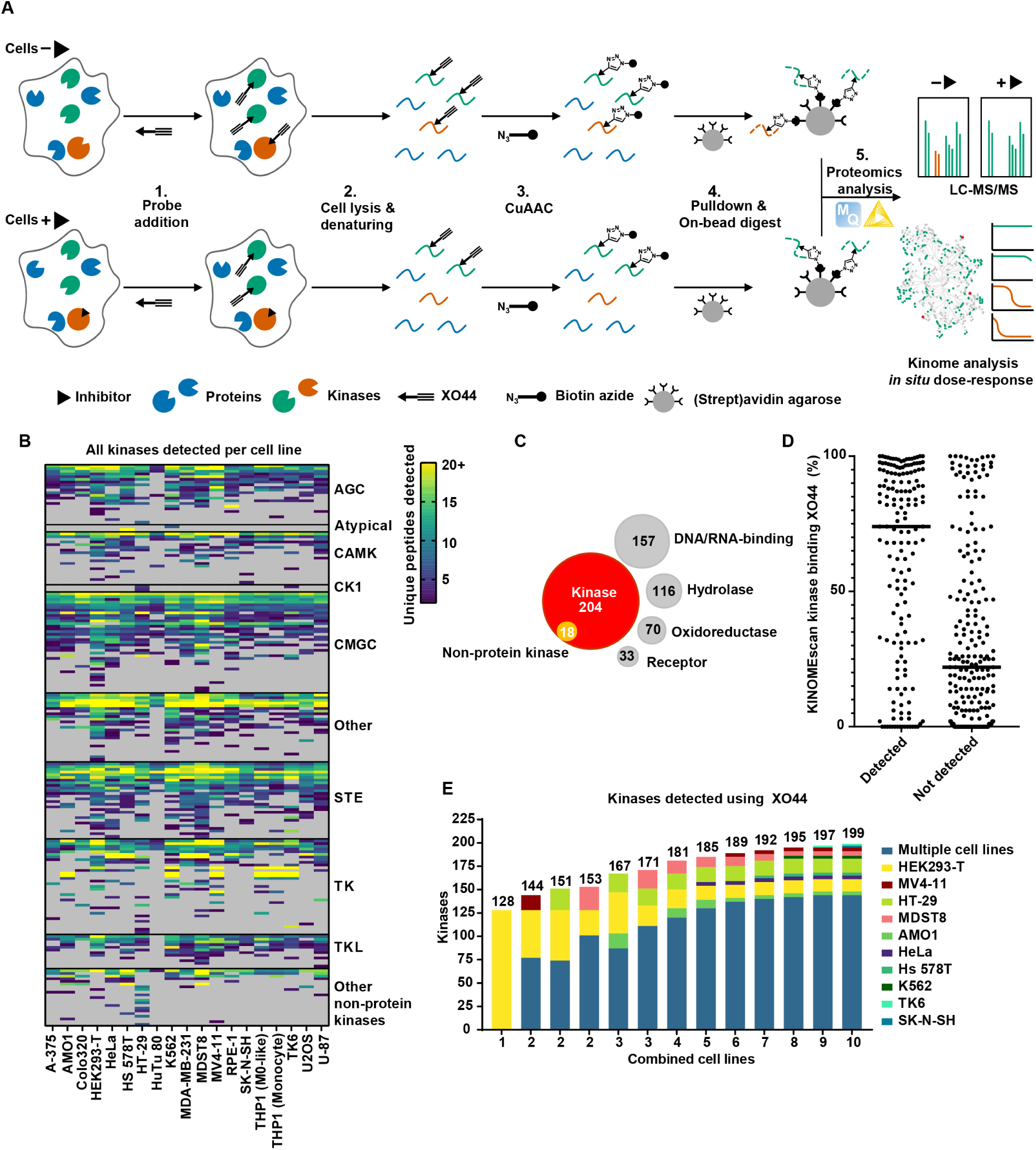
Overview chemical proteomics workflow and identification of optimal cell lines. **(A)** Graphical representation of competitive chemical proteomics workflow. **(B)** Unique peptides detected of probe-enriched kinases per cell line tested. Proteomics data is from n = 3-4 biological replicates of 1 µM XO44. **(C)** Molecular function analysis of all probe-enriched proteins from all tested cell lines. Main molecular functions are displayed. **(D)** Kinase binding of 1 µM XO44 as determined by KINOMEscan assay (Eurofins). Lines represent median binding values per condition. **(E)** Kinaseblender^37^ analysis of kinome coverage when combining data from different cell lines. See also Table S1.

### Selection of cell lines for optimal kinome coverage

Next, we aimed to evaluate the kinome coverage of XO44 across various cell lines to identify the optimal cell lines for broad coverage and to pinpoint kinases that are sufficiently abundant to be detected only in certain cell lines. To this end, 19 cell lines of diverse origins, such as myeloid cells (K562, MV4-11, THP-1, TK6), intestinal cells (Colo320-HSR, HT-29, MDST8, HuTu-80), brain cells (SK-N-SH, U87), and others (A375, AMO1, HEK293T, HeLa, Hs 578T, MDA-MB-231, RPE1 and U2OS) were treated with 1 µM XO44 or vehicle and processed for chemical proteomics analysis. Notably, for each cell line ∼ 0.75 mg of protein after lysis was used as starting point for the chemical proteomics. Label-free quantification (LFQ) in MaxQuant was used for protein quantification^36^ with the following criteria to define a kinase as a probe target: at least two unique peptides with an LFQ intensity ratio of at least two between probe- and vehicle-treated conditions, and appearance on a reference Uniprot kinase list (Keyword: KW-0418, Organism: 9606, Reviewed; Table S1).^2^ In total 204 distinct probe-enriched kinases were identified across all cell lines (Figure 1B, Table S1). 45 kinases were only identified in one cell line. The majority (37%) of the probe-enriched proteins were annotated with the keyword “kinase”, while the largest alternatives were DNA- or RNA-binding, hydrolase, oxidoreductase, or receptor (Figure 1C). To determine the binding affinity of XO44 to individual kinases, a biochemical KINOMEscan screen was performed (Figure 1D, Table S2). The results showed that the majority of kinases detected by the probe in the cellular screen showed also high binding affinity for XO44, with a median binding of 75%. In contrast, kinases not found in the chemical proteomics workflow generally showed lower binding, with a median binding of 21%. A Kinaseblender analysis was performed to investigate the cell lines with the highest complementarity in regard to kinome coverage.^37^ The cell line with the highest coverage was HEK293T (128 kinases). Combining it with a MV4-11 increased the coverage by 16-25 kinases (Figure 1E), with diminishing returns observed as the number of cell lines increased further. The combination of HEK293T and MV4-11 was therefore selected for optimal coverage and ease of handling.

### Increasing coverage of low abundant kinases

Next, we determined if the coverage of the kinome could be improved during the data analysis stage in MaxQuant. While label-free quantification allows for comparison across multiple samples, it has drawbacks, such as the need for consistent quantification and identification across several LC-MS runs. This can result in the inability to identify a peptide if its MS/MS spectra are of low quality or absent, leading to challenges in quantifying it. To address this issue, Cox and colleagues introduced the “Match Between Runs” (MBR) function, which transfers peptide identification across similar samples by matching m/z and retention time.^38,39^ This approach reduces missing intensities and enables the use of “peptide libraries” — data from samples included specifically to enhance peak identification. Although increasing the probe concentration for all samples could raise the number of labeled kinases, it would result in an underestimation of target engagement in competition experiments (Figure S4A). To facilitate peptide and kinase identification without compromising target engagement readouts, we created an experiment-specific “peptide library” (Figure 2A). This method involved processing MV4-11 cells with a high XO44 concentration (10 µM) in the absence of an inhibitor, alongside samples with varying inhibitor concentrations and a low probe concentration (1 µM). This approach allowed for the identification of peptides in samples with high concentration of probe, which could then be transferred to other samples, thereby increasing the efficiency of identification from 137 to 179 kinases (Figure 2B, Table S3) and increased peptide coverage for many others (Figure S4B). The kinases found only after matching were mostly low abundant, indicating poor engagement by 1 µM XO44 (Figure 2C). The reliability of these quantifications was confirmed by plotting the intensities of peptides identified by matching across replicates, which showed excellent correlation (Figure 2D).

**Figure 2:**
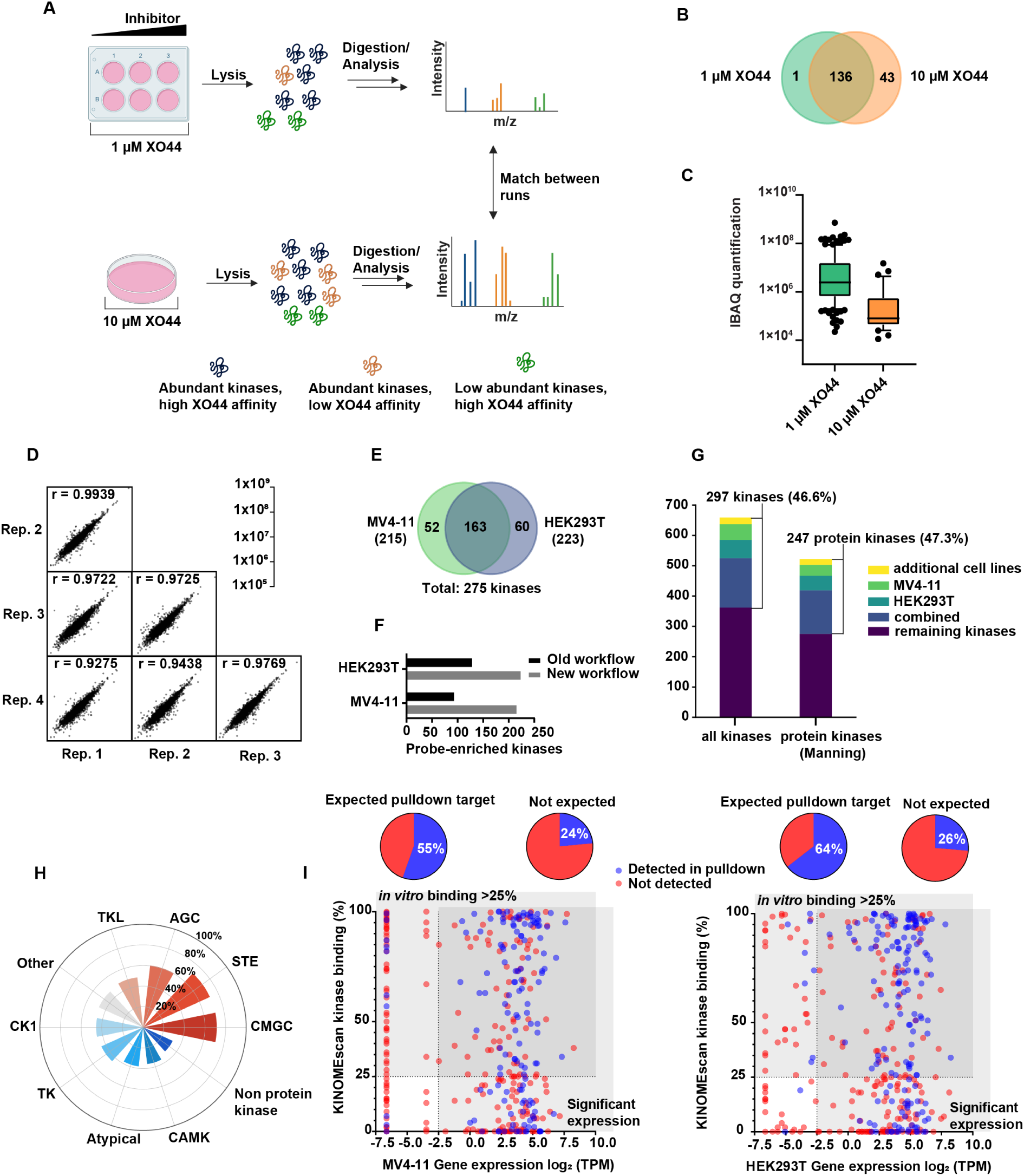
Peptide library increases kinome coverage and optimized workflow overview. **(A)** Graphical representation of the peptide library. Creation of an experiment-specific peptide library, involving samples processed with high probe concentration (10 µM) without inhibitors and samples with varying inhibitor concentrations and low probe concentration (1 µM), enabling efficient peptide identification and enhanced kinase identification through the ‘Match between runs’ function. **(B)** Probe-enriched kinases detected with high and low probe concentration. Increasing probe dose results in more detected kinases, indicating that at low probe dose these kinases are not efficiently engaged. Proteomics data is from n = 2-4 biological replicates. **(C)** iBAQ quantification of probe-enriched kinases indicates kinases quantified only after matching with a peptide library have low abundance. **(D)** Comparing the intensities of matched peptides between replicates by Pearson correlation indicates that matching with a peptide library results in reliable quantification results. See also Table S3. **(E, F)** Comparing the results of pulldown experiments on HEK293T and MV4-11 cells using the old and new workflow indicates a major improvement in kinome coverage with the same input material, resulting in 275 probe-enriched kinases in a single experiment. **(G)** Kinase coverage is around ∼ 50% across all kinases (protein kinases and metabolite kinases; Keyword: KW-0418, Organism: 9606, Reviewed)^2^ and Manning^1^ kinases. **(H)** Illustration of kinase coverage across the different kinase families. **(I)** Comparing biochemical binding of XO44 determined by KINOMEscan assay (Table S2) and mRNA levels of kinases from the FANTOM5 dataset^40^ creates overlap of kinases which are expected to be detected in a pulldown. Of those expected kinases, 55-64% are actually detected experimentally. See also Table S5 and S6.

Finally, we optimized the loading and type of beads to reduce the streptavidin background in order to detect low abundant kinases. Using streptavidin agarose and reducing bead loading improved sequence coverage and reduced ion suppression. This refined protocol also streamlined the pulldown workflow, allowing simultaneous preparation of multiple samples (Figure S5, Table S4). We tested the optimized method in MV4-11 and HEK 293T cell lines. A total of 275 distinct kinases were detected, with 215 identified in MV4-11 cells and 223 in HEK293T cells (Figure 2E, Table S5 and S6). The optimized workflow demonstrated an average two-fold increase in kinome coverage compared to the non-optimized workflow, using the same protein input of approximately 0.75 mg (Figure 2F). Combined with the number of kinases identified in the other cell lines of the panel screen (Table S1), a total of 297 kinases were identified, representing nearly half of the kinome, including both protein and metabolite kinases (Figure 2G). A similar coverage was observed when considering only the protein kinases, as defined by Manning.^1^ XO44 exhibited varying coverage across different kinase families. The homologs of yeast sterile kinases (STE) and CMGC (named after: cyclin-dependent kinases (CDK), mitogen-activated protein kinases (MAPK), glycogen synthase kinases (GSK), and CDK-like kinases), families were the best represented, with over 70% coverage (Figure 2H). In contrast, the non-protein kinases, as well as the calcium/calmodulin-dependent protein kinase (CAMK) and atypical families, were underrepresented with less than 40% coverage. This may be due to the probe’s lack of affinity for these groups’ structurally unique members.

To investigate the limitations of the current kinome coverage by XO44, its binding affinities to purified kinases as measured in the KINOMEscan assay and reported mRNA expression levels^40^ of HEK293T and MV4-11 were compared to the target engagement in the chemical proteomics experiment (Figure 2I). We could detect 55-64% of the kinases when we considered an *in vitro* binding of > 25% and a gene expression level of > -2.5 log2 transcript per million (TPM). This suggests that other probes with different scaffolds compared to XO44 may increase kinome coverage. Furthermore, kinases with low or no mRNA expression levels were generally not identified in our chemical proteomics workflow, which indicates that additional cell lines or conditions need to be screened to detect those kinases. Of interest, 36-45% of the kinases targeted by XO44 in the biochemical assay were expressed in the cell lines, but not detected in the chemical proteomics workflow. This suggests that the active site of those kinases was not available to the probe in the cellular context. This might be due to protein-protein interactions, post-translational modifications, high local substrate concentrations or the kinases were in an inactive conformation.

### Automated data processing and validation of the workflow using FDA-approved inhibitors

Next, we developed a data processing workflow using the KNIME Analytics Platform to automate the quantification of target engagement and selectivity profile of kinase inhibitors. The workflow requires the MaxQuant output proteingroups.txt file, a sample list, and a list of the proteins of interest, in this case Uniprot kinases [KW-0418].^2^ The workflow imputes missing data from a Gaussian distribution^41^ and filters for reliable quantification, appearance on the proteins of interest list, and probe-enrichment. The engagement of inhibitors is determined, and a pIC_50_ curve is fitted through the normalized LFQ values for the proteins of interest.

To validate our optimized chemical proteomics workflow with automated data processing, we determined the selectivity profile of the FDA-approved, reversible kinase inhibitor midostaurin, which is used for the treatment of acute myeloid leukemia. Target engagement was determined in a full dose-response manner across six concentrations, ranging from 0.1 nM to 10 µM. Each concentration was measured in biological duplicates, with the positive control and peptide library measured in biological quadruplicates and the negative control in biological duplicates. This setup enabled the generation of full dose-response curves for each engaged and enriched kinase. Competition of kinase labeling by midostaurin was analyzed and visualized in a radar plot, grouping kinases based on their families (Figure 3A). Midostaurin interacted with 50 kinases out of 275 enriched kinases in HEK293T and MV4-11 cells combined (Table S5 and S6). Midostaurin had a pIC_50_ of 8.15 ± 0.16, on its main target FLT3 in MV4-11 cells, which is in line with its inhibitory potency in a biochemical assay. Across the kinome, pIC_50_ values calculated from both cell lines showed a strong correlation with the exception of Aurora kinase A (Figure 3B). The engagement of Aurora kinase A by midostaurin was found to be 100-fold lower in HEK 293T cells, potentially due to differences in phosphorylation state.^42^ Eighteen targets of midostaurin identified using our method were not inhibited in the kinobeads assay, most of which had a pIC_50_ value lower than 6, highlighting the sensitivity of our workflow (Figure 3C). To validate a potential off-target of midostaurin, we chose SIK2 as a representative example. To this end, we compared the SIK2 engagement of midostaurin across various methods, including RapidFire, a biochemical activity-based assay using the full length protein, and NanoBRET (Figure 3D). As expected, the biochemical pIC_50_ was significantly higher than the cellular pIC_50_, but the pIC_50_ value in the NanoBRET was comparable to our chemical proteomics readout.

**Figure 3.**
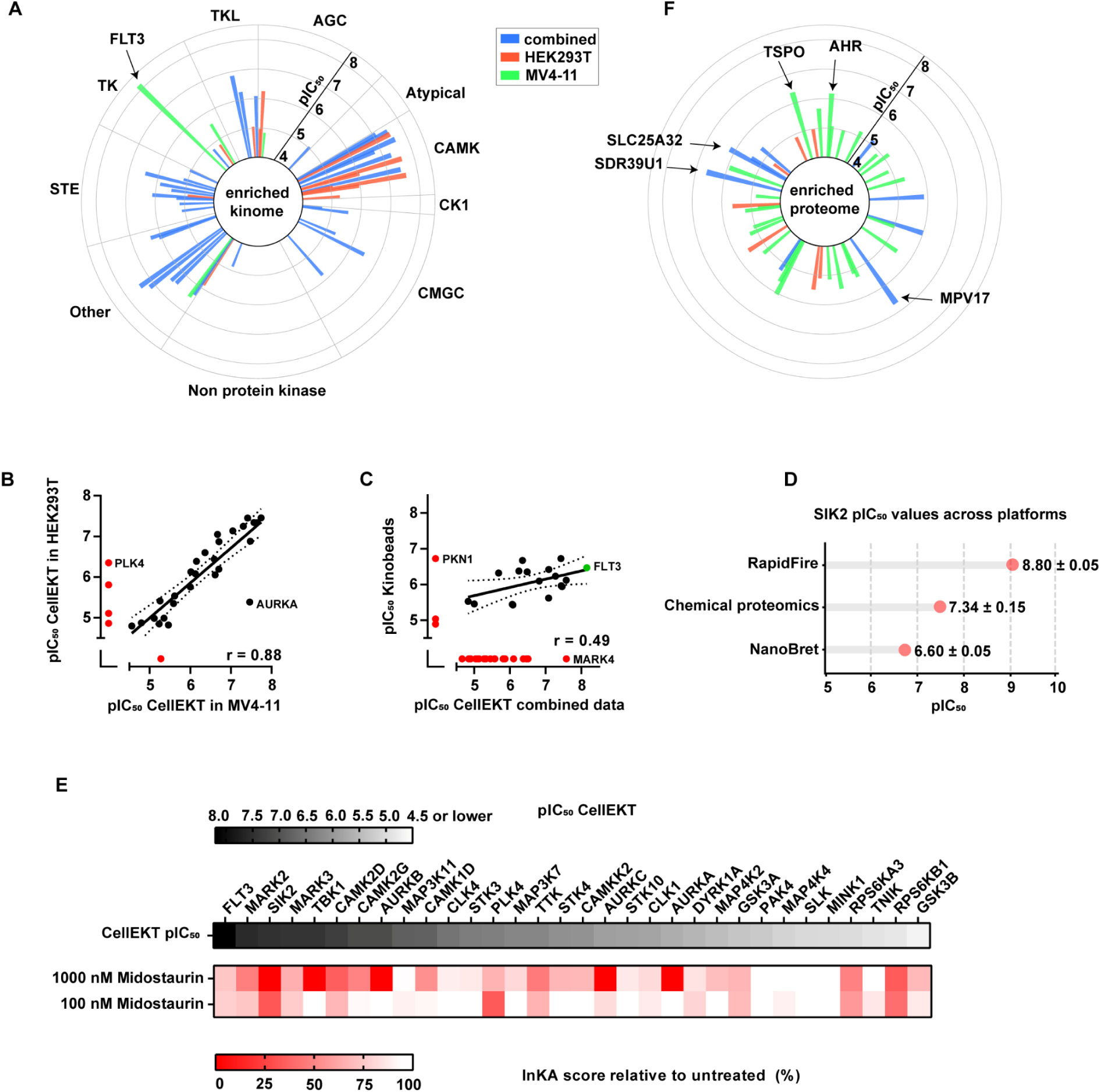
Testing midostaurin in the CellEKT assay. **(A)** Radar plot depicting the kinase targets and binding affinities of midostaurin identified with a good logarithmic fit (Table S5 and S6). Each spike represents one target which was measured at 6 different concentrations from 0.1 nM to 10,000 nM and the data is presented as mean values of n = 2 biological replicates. The length of each spike indicates the determined pIC_50_ and the color in which cell line HEK293T (red), MV4-11 (green) or both cell lines (blue), target occupancy was determined. If target occupancy was determined in both cell lines the pIC_50_ values were averaged. **(B)** Correlation of kinase pIC_50_ of midostaurin between MV4-11 and HEK293T in CellEKT assay. Kinases which were found not inhibited are marked in red. **(C)** Correlation of kinase pIC_50_ values of midostaurin between XO44 chemical proteomics and kinobeads assay.^9^ Kinases which were found not inhibited are marked in red. **(D)** Comparison of determined pIC_50_ values of midostaurin on SIK2 using the RapidFire, chemical proteomics, and NanoBRET platforms. **(E)** Comparison of determined pIC_50_ values using CellEKT, as determined in a full dose response curve, with the effects on kinase activity and kinase substrates (INKAscore)^43^ as determined by phosphoproteomics (Table S9), conducted in n = 2 biological duplicates for each concentration. **(F)** Radar plot illustrating the targets and binding affinities of midostaurin across the enriched proteome, with the kinome excluded.

Furthermore, to correlate the target engagement profile with modulation of target kinase activity, we performed a phosphoproteomics experiment in MV4-11 cells using two concentrations of midostaurin (Figure 3E). Gratifyingly, the Integrative Kinase Activity (InKA) score^43^ from the phosphoproteomics data (Table S9) correlated well with the pIC_50_ values across the 32 targets determined by our chemical proteomics workflow. Finally, we identified also several non-kinase targets engaged by midostaurin in both cell lines (Figure 3F, S6A). For example, a dose-dependent inhibition of the Aryl hydrocarbon receptor (AHR) (Figure S6B) was observed. This is in line with the previous report that midostaurin and its active metabolites are inducers of CYP1A2, an effect mediated by AHR.^44^

To further validate our chemical proteomics workflow, we also profiled three FDA-approved covalent, irreversible BTK inhibitors (ibrutinib, acalabrutinib and zanubrutinib) used for the treatment of B-lymphocyte tumors (Figure 4A and 4B).^45^ All three BTK inhibitors were screened in HEK293T (Table S7) and MV4-11 (Table S8). Ibrutinib engaged 25 kinases, of which 10 were detected in both cell lines (Figure 4C), with a pIC_50_ ranging from of 8.6 (BTK) to below 5 (CDK14, STK38L, LIMK1 and MAPK13). Notably, 14 kinases were identified with pIC_50_ values below 6. Zanubrutinib engaged 16 kinases and acalabrutinib showed the least off targets with TEC, RIPK2 and TGFBR1. To validate the observed off-targets, we measured the activity of the three BTK inhibitors in a KINOMEscan (Table S2) and compared to the % inhibition to the chemical proteomics profiled. All engaged targets were confirmed with a general trend of reduced activity, which aligns with previous findings comparing biochemical assays to cellular activity data.^24^ Of note, acalabrutinib did not show any selectivity over TEC, which is an off-target associated with an increased bleeding risk (Figure 4B).^46^

**Figure 4.**
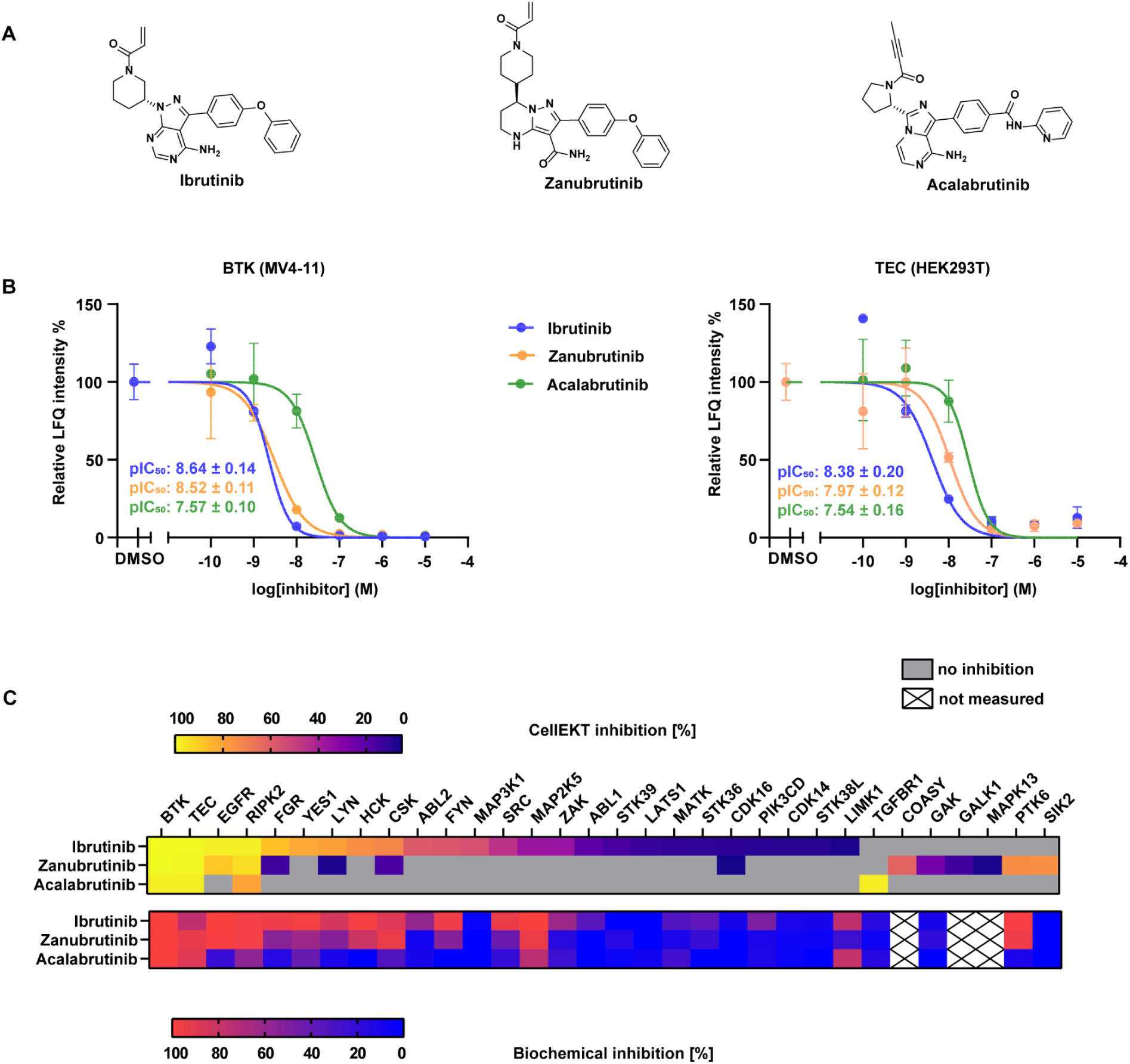
Comparing BTK inhibitors in the CellEKT assay. **(A)** Chemical structures of ibrutinib, zanubrutinib and acalabrutinib. **(B)** Full dose response curve of TEC (Table S7) as determined in HEK293T and of BTK as determined in MV4-11 cells (Table S8) for ibrutinib, zanubrutinib and acalabrutinib. Target engagement is determined at 6 different concentrations from 0.1 nM to 10,000 nM and data is presented as mean values ± SEM of n = 2 biological replicates. **(C)** Comparison of kinase % inhibition by CellEKT (Table S7 and S8) and % inhibition at 1 µM inhibitor concentration determined by KINOMEscan assay (Table S2). For the CellEKT analysis, if a pIC_50_ was determined in both cell lines, the values were averaged. The determined pIC_50_ values measured by CellEKT were transformed to % inhibition by: % Inhibition=([I]/[I]+10^−pIC^^50^) ×100 with inhibitor concentration [I] = 1 µM to match the concentration at which the KINOMEscan assay was performed.

### Additional probes ALX005 and ALX011

Our validated chemical proteomics workflow detected 275 kinases, but as indicated above (Figure 2H) we missed kinases mainly from the TK, atypical, CAMK and non protein kinase families. To address this gap, we decided to design two probes with a different scaffold from that of XO44. To this end, we converted a previously reported promiscuous kinase inhibitor^47^ into two broad-spectrum probes ALX005 and ALX011 (Figure 5A). Using a KINOMEscan (Table S2), we confirmed that the probes had broad-spectrum kinase activity and inhibited 51 additional kinases compared to XO44 (Figure 5B). Subsequently, we tested these probes in HEK293T and MV4-11 cells at concentrations of 1 µM and 10 µM (Figure 5C, Table S10). In total 304 kinases were enriched across HEK293T and MV4-11. Of note, ALX005 and ALX011 enriched for additional TK members such as the ephrin receptors A and B (EphB1-B3, EphA2-A4 and A7).

**Figure 5.**
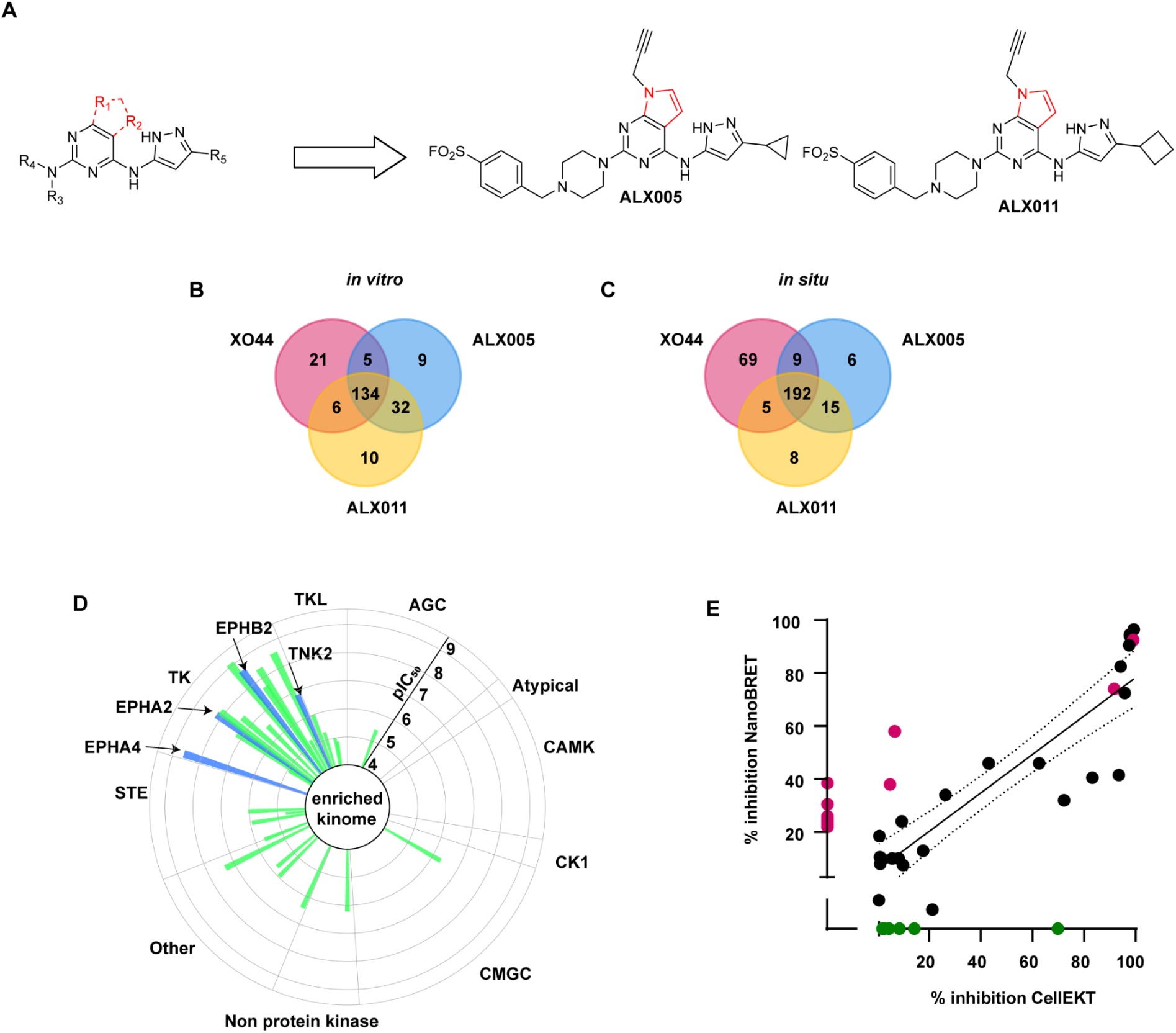
Profiling of ALX005/ALX011 and kinome-wide analysis using a probe cocktail to evaluate the selectivity profile of dasatinib. **(A)** Chemical structures of ALX005 and ALX011 derived from pyrrolo-pyrimidine core and XO44. **(B)** Shared and unique kinases with ≥ 50% inhibition at 1 µM as determined by KINOMEscan assay (Table S2). **(C)** Shared and enriched kinases across MV4-11 and HEK293T as determined by CellEKT (Table S10). **(D)** Radar plot depicting the kinase targets and binding affinities of dasatinib determined in MV4-11 identified with a good logarithmic fit (Table S11). Each spike represents one target which was measured at 6 different concentrations from 0.1 nM to 10,000 nM and the data is presented as mean values of n = 2 biological replicates. The length of each spike indicates the determined pIC_50_. The targets identified exclusively by ALX005 and/or ALX011, as depicted in Figure C, and demonstrating target occupancy with dasatinib are highlighted in blue. **(E)** Correlation plot comparing the percentage inhibition determined by NanoBRET^15^ with the percentage inhibition determined by CellEKT. The target occupancy of dasatinib was determined at 100 nM with NanoBRET. The determined pIC_50_ values measured by CellEKT were transformed to % inhibition by: % Inhibition=([I]/[I]+10^−pIC^^50^) ×100 with inhibitor concentration [I] = 100 nM. Kinases which were found inhibited across both platforms are illustrated in black with R^2^ = 0.81. Kinases that were found inhibited by CellEKT but were not analyzed by NanoBRET are illustrated in green. Kinases measured by NanoBRET that are not inhibited in the CellEKT assay, or have a poor logarithmic fit, are illustrated in red.

To exemplify the expanded kinome coverage, we determined the target engagement profile of the FDA-approved tyrosine kinase inhibitor dasatinib using a probe cocktail of XO44, ALX005 and ALX011 in MV4-11 cells. In this set up we detected 29 targets of dasatinib which were engaged with pIC_50_ ranging from 9.1 (EPHB4) to 4.7 (MAP4K3) (Figure 5D, Table S11). The chemical proteomics profile correlated well (R^2^ = 0.81) with the percentage inhibition at 100 nM dasatininb measured using NanoBRET^15^ (Figure 5E). Notably, the tyrosine kinases TNK2, EPHA2, EPHA4 and EPHB2, specifically enriched by the ALX-probes were previously not detected as off-targets by XO44. This demonstrated the complementarity of ALX005 and ALX011 to XO44.

### Identification of the kinase substrate profiles

Our optimized chemical proteomics workflow expanded the coverage of the kinome. Consequently, we hypothesized that a broader scope of phosphosites (peptide substrates) will be covered by the kinases detected in our platform. To quantify this, we compared the kinases with those for which substrate preferences have been recently studied.^30,48^ Among 392 unique kinases with well-characterized substrate motifs (303 serine/threonine kinases, 93 tyrosine kinases, with four kinases with dual-specificity), 222 protein kinases are enriched by our probes (176 serine/threonine and 46 tyrosine kinases). We also identified 82 enriched kinases for which substrate motifs are not yet established (Figure 6A).

**Figure 6.**
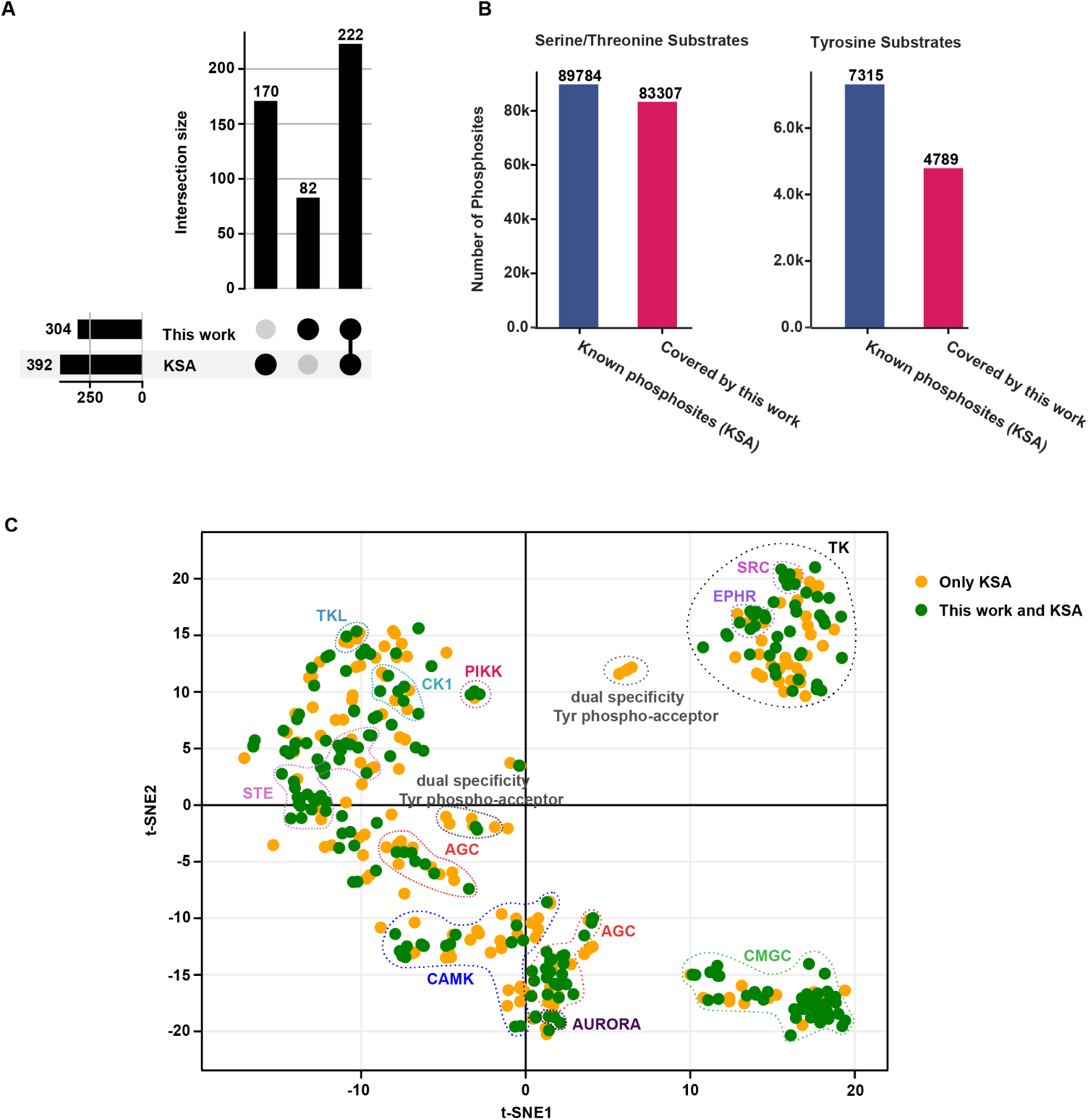
Expansion of the kinase-substrate coverage. **(A)** Upset plot showing kinases covered by this work compared to kinase-substrate atlas (KSA).^30,48^ The upper bar plot shows the intersection sizes, highlighting the number of kinases uniquely identified in this work, those covered by both, and those exclusive to KSA. The lower plot presents the overall counts of kinases covered by each study and their overlap. **(B)** Coverage of known serine/threonine and tyrosine phosphosites from high^49^ and low-throughput^50^ approaches. A substrate was considered confidently covered by a kinase if the kinase was found in the 90^th^ percentile according to the scoring and percentile ranking from kinase-substrate atlas^30,48^ **(C)** t-SNE plot of kinases clustered by their amino-acid sequence preference. Each dot represents a kinase, colored orange if only covered by KSA, and green if covered by both this work and KSA. Known families of kinases are highlighted with dashed circles and labeled using reference colors from the phylogenetic tree of the human kinome.

We assessed the substrate profiles of the 222 kinases with well-characterized substrate motifs, including all human phosphosites that have been demonstrated as substrates of these kinases in low-throughput and high-throughput studies.^29,30,49,50^ Kinases engaged by our platform demonstrated a substantial coverage of the human kinase targetome (Figure 6B), including 83,307 unique serine/threonine-containing substrates (92.7% of the known Ser/Thr kinase substrates) and 4,789 tyrosine-containing substrates (65.4% of the known Tyr kinase substrates).

Moreover, we performed clustering analysis to human kinases included in the kinase-substrate atlas (KSA) based on their preferred amino-acid sequences around the phosphorylation site, and plot the kinases in a t-SNE plot by their motif similarity in Figure 6C. In the t-SNE plot, kinases with similar substrate targets are closer to each other. The 222 kinases enriched by our chemical proteomics workflow, marked in green on the t-SNE plot, show a broad coverage of the kinome (Figure 6C).

Taken together, these observations suggests that our workflow is able to detect the majority of kinases, which are able to phosphorylate the majority of known protein substrates. It is conceivable that the kinases profiled by our platform are probably responsible for a large proportion of biological processes regulated by kinase-mediated signaling. This expansion provides valuable insights into kinase functions and their roles in various biological processes.

## Discussion

Determination of cellular target engagement and selectivity profiles of kinase inhibitors is crucial to understand their mode-of-action and to prevent adverse effects. Chemical proteomics, specifically using the XO44 probe, has advanced the study of the interaction landscape of kinase inhibitors in native biological contexts. XO44 was able to detect 133 kinases^25^, which represents 21% of the total kinome. Thus, there is a need to increase the kinase coverage and to develop a robust protocol to assess the target engagement of inhibitor across the total kinome.

To address this question, we optimized the sample preparation and chemical proteomics workflow to enhance detection and analysis of kinase targets. Key optimizations included refining cell density, cell lysis conditions, and ligation efficiency. Fluorescence and gel-based techniques helped track protein labeling and adjust pull-down conditions for better results. Testing across various cell lines revealed that HEK293T and MV4-11 provided the best kinome coverage. Enhancements in data processing, such as the “Match Between Runs” function, significantly increased kinase identification, particularly for low-abundance targets. Optimizing bead loading further improved detection sensitivity. Profiling using FDA-approved inhibitors validated the accuracy of the workflow, with midostaurin and three BTK inhibitors demonstrating expected target engagement and revealing novel off-target interactions. Furthermore, two additional probes ALX005 and ALX011 were developed, enriching additional kinases from the tyrosine kinase subfamily and others. In total we could detect 304 kinases (48% of kinome) with our optimized chemical proteomics workflow, which we termed CellEKT (Cellular Endogenous Kinase Targeting). This expanded kinome coverage allows for more comprehensive substrate profiling, providing insights into kinase-mediated biological processes and potential off-target effects of kinase inhibitors. In a longer perspective this might allow for identification of signature profiles based on more relevant quantitative inhibition and selectivity data in living cells. Thereby improving prediction and reducing major off target kinase liabilities such as cardiotoxicity, bone marrow toxicity and genetic instability caused by disruption of kinase signaling.

Biochemical data may fail to predict inhibitor behavior in cells^51^ and often overestimates potency, as we have confirmed for SIK2 inhibition by midostaurin (Figure 3D). This discrepancy was also observed with lysate-based Kinobead readouts, highlighting the importance of assessing cellular inhibitor selectivity profiles.^15,24^ Techniques like CellEKT, NanoBRET and CETSA are capable of such assessments.

CETSA is a powerful tool for full proteome drug target screening in complex biological systems but misses targets that are not CETSA-compatible and provides limited selectivity information within enzyme families such as the kinome.^16^ NanoBRET allows thorough evaluation of cellular kinase selectivity at thermodynamic equilibrium but is limited in flexibility for broad kinome selectivity profiling across different cell lines or in complex, native biological contexts. Conversely, CellEKT, despite its limitation in assessing target engagement at thermodynamic equilibrium due to the covalent nature of the probe, enables the IC50 determination of an inhibitor across 304 kinases and can be applied to any cell line, primary cell, and organoid. CellEKT assesses the selectivity of a compound fully automatically across half of the kinome and other non-kinase proteins. Thus, CellEKT offers the possibility to determine cellular target engagement in relevant biological settings and to couple it to phenotypic or target modulations assays, such as phosphoproteomics.

Although we did not study specifically weaker affinity binders, our platform was able to capture many low-affinity interactions that correlated well with NanoBRET data. When it comes to fast k_off_/slow k_on_ kinetic binders, CellEKT should be used with caution, and the probe incubation time or concentration needs to be optimized prior to the experiment. Moving forward, CellEKT can be further improved to expand its kinome coverage. Stimulating or synchronizing cells should lead to detectable expression levels of pulsatile expressed kinases.^52^ Additionally, increased coverage could be achieved using different mass spectrometry acquisition workflows, such as Data Independent Acquisition (DIA)^53^, and novel complementary probes.

## Conclusion

The CellEKT platform with an optimized XO44 chemical proteomics workflow significantly improves the profiling of kinase inhibitor interactions within native cellular environments. CellEKT expands the kinome coverage to 48% and its readout is validated by orthogonal biochemical assays, NanoBRET, and phosphoproteomics. A robust and versatile cellular kinase screening platform, CellEKT can be applied to various biological contexts. Future advancements, including novel mass spectrometry methods and complementary probes, are anticipated to further expand kinome coverage and enhance CellEKT’s role in supporting drug discovery and development.

## Significance

Kinases are enzymes that phosphorylate proteins or cellular metabolites, playing a key role in regulating protein function, signaling pathways, and cellular processes. They are important drug targets, particularly in the treatment of cancer and autoimmune diseases. As a result, the discovery of kinase inhibitors is a major focus in both academic and industrial research, with over 80 FDA-approved kinase inhibitors available. However, many of these inhibitors target the conserved ATP-binding site in kinases, often leading to off-target interactions and adverse side effects. Due to issues related to toxicity and efficacy, most kinase inhibitors fail in clinical trials, making it essential to understand their interactions with targets, especially for non-oncological applications.

An important aspect of drug development is determining the concentration needed to fully engage the target without causing undesirable off-target effects in a cellular environment. In this study, we introduce CellEKT (Cellular Endogenous Kinase Targeting), an optimized and robust chemical proteomics platform for investigating the cellular target engagement of endogenously expressed kinases. Using the sulfonyl fluoride-based probe XO44 and two newly developed probes, CellEKT expanded the coverage of the human kinome from 21% to 48%. This optimized workflow enabled the determination of the kinome interaction landscape of both covalent and non-covalent drugs across more than 300 kinases, expressed as half-maximal inhibitory concentrations (IC50), which were validated using platforms like phosphoproteomics and NanoBRET. With CellEKT, we linked target engagement profiles to their corresponding substrate space, providing insights into drug mechanisms. CellEKT is a powerful tool for decoding drug actions and guiding the discovery and development of new therapeutics.

## Supporting information

Table S1. Cell line kinome coverage of XO44

Table S2. Uniprot kinases and DiscoveRx KINOMEscan data

Table S3. Using high probe dose samples as peptide library

Table S4. Chemical proteomics method comparison

Table S5. KNIME output kinome selectivity assay-Midostaurin_HEK293T

Table S6. KNIME output kinome selectivity assay-Midostaurin_MV4-11

Table S7. KNIME output kinome selectivity assay-BTK_inhibitors_HEK293T

Table S8. KNIME output kinome selectivity assay-BTK_inhibitorsMV4-11

Table S9. phosphoproteomics_data_midostaurin

Table S10. Kinase coverage ALX005_ALX011

Table S11. Probe cocktail

## Supplemental information

Supplemental Information can be found online at [LINK]

## Acknowledgments

The authors would like to thank the laboratory of Henk M.W. Verheul at the Radboud University Medical Center for kindly providing the HT-29 cell line, as well as laboratory of Christoph Driessen for kindly providing the AMO1 cell line. Financial support for this work was provided by F. Hoffmann-La Roche through the pRED ROADS program and the Oncode Institute. Further, support for this publication was received by Oncode Accelerator, a Dutch National Growth Fund project under grant number NGFOP2201

## Author contributions

J.R. performed experiments, collected and analyzed data, performed the chemical proteomics analysis on the timsTOF and contributed to the writing of the manuscript. B.G. performed experiments, collected and analyzed data, performed the phosphoproteomics analysis on the timsTOF and contributed to the writing of the manuscript. M.B. and M.P. performed experiments, collected and analyzed data. A.L. performed the synthesis. A.V and J.D.Z. performed the computational study and contributed to the writing of the manuscript. A.P.A.J. conceptualized and wrote the KNIME data analysis workflow. B.I.F. performed the proteomics analysis on the Q Exactive. P.W. and B.W. developed the RF assay and A.T. performed the experiment and analyzed the data. H.W. developed the NanoBRET assay and R.H. performed the experiment and analyzed the data. J.P.M. and P.G.B. provided material for the cell line study. A.C.R., M.S., S.K. and U.G. conceived the project, and contributed to the writing of the manuscript.

## Supporting citations

The following references appear in the supplemental information: ^54,55^

## RESOURCE AVAILABILITY

### Lead contact

Further information and requests for resources and reagents should be directed to and will be fulfilled by the lead contact, Mario van der Stelt.

### Materials availability

ALX005 and ALX011 are available upon request.

### Data and code availability

The raw (phospho)proteomics datasets generated during this study are available at PRIDE accessions: PXD035544, PXD035540, PXD035542, PXD035549, PXD035619, PXD035550, PXD035551, PXD054026, PXD053911 and PXD055387 will be publicly available as of the date of publication. The accession numbers for these data as well as existing, publicly available data are listed in the key resources table.

The code necessary to run the automated KNIME analysis workflow along with instructions can be found at DOI: https://doi.org/10.5281/zenodo.7656526

Any additional information required to reanalyze the data reported in this paper is available from the lead contact upon request.

## EXPERIMENTAL MODEL AND SUBJECT DETAILS

### Cell culture

Colo320-HSR (Female human origin) and MDST8 (Gender undetermined) cells were cultured at 37 °C under 7% CO_2_ in in RPMI 1640 (R5886, Merck) with GlutaMAX, 1% glucose (G8769, Merck), 10% FCS (Thermo Fisher), 1 mM sodium pyruvate, and penicillin and streptomycin (200 µg/mL each, Duchefa). Cells were passaged twice a week at 80-90% confluence by trypsinization.

HeLa (Female human origin) and U2OS (Female human origin) cells were cultured at 37 °C under 7% CO_2_ in DMEM (D6546, Merck) with GlutaMAX, 10% NBS (Thermo Fisher), and penicillin and streptomycin (200 µg/mL each, Duchefa). Cells were passaged twice a week at 80-90% confluence by trypsinization.

SK-N-SH (Female human origin) and U-87 (Male human origin) cells were cultured at 37 °C under 7% CO_2_ in in in DMEM (D6546, Merck) with GlutaMAX, 10% FCS (Thermo Fisher), and penicillin and streptomycin (200 µg/mL each, Duchefa). Cells were passaged twice a week at 80-90% confluence by trypsinization.

Hs 578T (Female human origin) and MDA-MB-231 (Female human origin) cells were cultured at 37 °C under 7% CO_2_ in in RPMI 1640 (R5886, Merck) with GlutaMAX, 10% FCS (Thermo Fisher), and penicillin and streptomycin (200 µg/mL each, Duchefa). Cells were passaged twice a week at 80-90% confluence by trypsinization.

RPE-1 (Female human origin) cells were cultured at 37 °C under 7% CO_2_ in in DMEM/F12 (D8062, Merck) with GlutaMAX, 10% FCS (Thermo Fisher), and penicillin and streptomycin (50 µg/mL each, Duchefa). Cells were passaged twice a week at 80-90% confluence by trypsinization.

HT-29 (Female human origin) cells were cultured at 37 °C under 7% CO_2_ in in DMEM (D6546, Merck) with GlutaMAX, 5% FCS (Thermo Fisher), 10 mM HEPES, and penicillin and streptomycin (200 µg/mL each, Duchefa). Cells were passaged twice a week at 80-90% confluence by trypsinization.

HuTu-80 (Male human origin) cells were cultured at 37 °C under 7% CO_2_ in in DMEM/F12 (D8062, Merck) with GlutaMAX, 10% FCS (Thermo Fisher), 10 mM HEPES, and penicillin and streptomycin (200 µg/mL each, Duchefa). Cells were passaged twice a week at 80-90% confluence by trypsinization.

HEK293T (Female human origin) cells were cultured at 37 °C under 7% CO_2_ in DMEM (D6546, Merck) with GlutaMAX, 10% NBS (Thermo Fisher), and penicillin and streptomycin (200 µg/mL each, Duchefa). Cells were passaged twice a week at 80-90% confluence by resuspension in fresh medium.

A375 (Female human origin), AMO1 (Female human origin), K562 (Female human origin) and THP-1 (Male human origin) cells were cultured at 37 °C under 7% CO_2_ in RPMI 1640 (R5886, Merck) with GlutaMAX, 10% FCS (Thermo Fisher), 1 mM sodium pyruvate, and penicillin and streptomycin (200 µg/mL each, Duchefa).

TK6 (Male human origin) cells were cultured at 37 °C under 7% CO_2_ in RPMI 1640 (R5886, Merck) with GlutaMAX, 10% horse serum (Thermo Fisher), 1 mM sodium pyruvate, and penicillin and streptomycin (200 µg/mL each, Duchefa).

MV4-11 (Male human origin) and SEM (Female human origin) cells were cultured at 37 °C under 5% CO_2_ in IMDM (I3390, Merck) with GlutaMAX, 10% FCS (Thermo Fisher), and penicillin and streptomycin (200 µg/mL each, Duchefa).

THP-1 cells were differentiated and stimulated by plating 3.0×10^6^ cells in 6 cm dishes in 5 mL complete RPMI 1640 containing 20 ng/mL Phorbol 12-myristate 13-acetate (PMA, Focus Biomolecules) for 24 h. Then, medium was replaced with serum-free RPMI 1640 with 0.1% delipidated BSA (Merck) with GlutaMAX, 1 mM sodium pyruvate, and penicillin and streptomycin (M0-like) for 24 h. Cell lines were tested regularly for mycoplasma contamination. Cultures were discarded after 2-3 months of use.

## METHOD DETAILS

### Cell treatment for chemical proteomics

Adherent cells were cultured in 6-well plates or 6 cm dishes (Sarstedt) and grown to 80-90% confluency, aiming to recover approximately 0.75 mg of total protein per condition after lysis. Prior to treatment, the growth medium was aspirated, and the cells were gently washed with 2 mL DPBS. Cells were treated with kinase inhibitor (1.11X) or vehicle (0.111% DMSO) in the corresponding serum-free culture medium supplemented with 0.1% delipidated BSA (Merck) and in case of IMDM or DMEM with additionally 10 mM HEPES (treatment medium). The kinase inhibitor was tested at six concentrations ranging from 0.1 nM to 10,000 nM in n = 2 biological replicates per concentration (12 samples total). Vehicle treatments were conducted across 10 samples, including: Negative control (n = 2), Positive control (n = 4) and Peptide library treatments (n = 4). Cells were incubated with the kinase inhibitor or vehicle at 37°C for 1 h. Following this, a solution of XO44 (10X for positive control and competition samples, or 100X for peptide library samples) or vehicle (1% DMSO or 0.2% DMSO final) was added in treatment medium. Cells were incubated at 37 °C for an additional 30 min. The medium was aspirated, and treatment was halted by the addition of 1 mL ice-cold DPBS. Cells were then scraped off on ice, collected into 1.5 mL Eppendorf tubes, and spun down again (1,000 *g*, 5 min, 4 °C). The supernatant was removed, and cell pellets were snap-frozen in liquid nitrogen and stored at -80 °C.

For cell line screening or probe profiling experiments, the initial 1 h incubation step was omitted, and cells were incubated with the probe (n = 3-4) or vehicle for 30 min directly before proceeding with collection.

Suspension cells in log phase were spun down (300 *g*, 5 min) and resuspended at 1.0×10^6^ cells per mL in the corresponding serum-free culture medium supplemented with 0.1% delipidated BSA (Merck) and in case of IMDM or DMEM, 10 mM HEPES. Amount of total cells was adjusted to receive around 0.75 mg of final protein after cell lysis. Cells were treated with kinase inhibitor (1.11X) or vehicle (0.111% DMSO). The kinase inhibitor was tested at six concentrations ranging from 0.1 nM to 10,000 nM in n = 2 biological replicates per concentration (12 samples total). Vehicle treatments were conducted across 10 samples, including: Negative control (n = 2), Positive control (n = 4) and Peptide library treatments (n = 4). Cells were incubated with the kinase inhibitor or vehicle at 37 °C for 1 h. Following this, a solution of XO44 (10X for positive control and competition samples, or 100X for peptide library samples) or vehicle (1% DMSO or 0.2% DMSO final) was added in treatment medium. Cells were incubated for an additional 25 min at 37 °C. Cells were spun down (300 *g*, 5 min, 37 °C), resuspended in ice-cold DBPS (1 mL), collected into 1.5 mL Eppendorf tubes, and spun down again (1,000 *g*, 5 min, 4 °C). The supernatant was removed, and cell pellets were snap-frozen in liquid nitrogen and stored at -80 °C.

For cell line screening or probe profiling experiments, the initial 1 h incubation step was omitted, and cells were incubated with the probe (n = 3-4) or vehicle for 30 min directly before proceeding with collection.

### Chemical proteomics assay for cell line screen

Screening the cell lines was performed with a method adapted from previous experience with chemical proteomics^34^ and usage of XO44 as kinase probe^25^ with modifications based on the gel-based results. Cell pellets were lysed completely by vortexing in 400 µL ice-cold lysis buffer containing 50 mM HEPES pH 7.5, 150 mM NaCl, 1 mM MgCl_2_, 0.1% (w/v) Triton X-100, 1X cOmplete™ EDTA-free Protease Inhibitor Cocktail (Roche) and 25 U/mL benzonase (sc-202391, Santa Cruz). Protein concentrations were measured by BCA assay (Thermo Fisher), equalized and 360 µL lysate was transferred to 2 mL Eppendorf tubes containing 40 µL 10% SDS (1% final), vortexed and left at room temperature.

The lysates were subjected to a click reaction for 1 h with freshly prepared click mix (31.2 µL per sample: 16 µL 25 mM CuSO_4_ in MilliQ, 3.2 µL 25 mM THPTA in DMSO, 4 µL 2.5 mM biotin-N_3_ in DMSO and 8 µL 50 mM TCEP.HCl freshly dissolved in DPBS). Proteins were precipitated by addition of HEPES buffer (80 µL, 50 mM HEPES pH 7.5), MeOH (666 µL), CHCl_3_ (166 µL) and MilliQ (150 µL), vortexing after each addition. After spinning down (1,500 *g*, 10 min) the upper and lower layer were aspirated and the protein pellet was resuspended in MeOH (600 µL) by sonication (Qsonica Q700 Microplate Sonicator, 2 × 10 s pulses, 10% amplitude). The protein was spun down (20,000 *g*, 5 min) and the supernatant was discarded. The proteins were redissolved in 6 M urea (500 µL) with 25 mM NH_4_HCO_3_ for 15 min, followed by reduction (65 °C, 15 min, 800 rpm shaking) with DTT (5 µL, 1 M in MilliQ). The samples were allowed to reach room temperature and proteins were alkylated (30 min) with IAA (40 µL, 0.5 M in MilliQ) in the dark. 140 µL SDS (10% w/v) was added and the samples were spun down (1,000 *g*, 5 min). The samples were then transferred to 15 mL Falcon tubes containing 5 mL PBS and 100 µL avidin agarose slurry (Pierce, 100 µL of a 50% slurry, prewashed twice with 6 mL PBS + 0.5% SDS and once with 6 mL PBS) and the samples were incubated for 2 h while rotating. The beads were then spun down (2,000 *g*, 2 min) and washed (3 x PBS + 0.5% SDS, 2 x PBS, 1 x MilliQ). The beads were resuspended in digestion buffer (250 µL, 100 mM Tris pH 7.8, 100 mM NaCl, 1 mM CaCl_2_, 2% (v/v) acetonitrile, sequencing grade trypsin (Promega, 0.25 µg)), transferred to low-binding tubes (Sarstedt) and incubated while shaking in Eppendorf tube shaker overnight (16 h, 37 °C, 1,000 rpm). Trypsin was quenched with 12.5 µL formic acid (ULC-MS grade) and the beads were removed by filtration using a Bio-Spin column (BioRad, 400 *g*, 5 min), collecting the flow-through in a new 2 mL tube. Peptides were desalted using C18 StageTips^56^ preconditioned with 50 µL MeOH, 50 µL of 0.5% (v/v) FA in 80% (v/v) acetonitrile/MilliQ (solution B) and 50 µL 0.5% (v/v) FA in MilliQ (solution A) by centrifugation (600 *g*, 1-2 min). The peptides on the StageTip were washed with solution A (100 µL) and eluted into new low-binding tubes using solution B (100 µL). Samples were concentrated using an Eppendorf speedvac (Eppendorf Concentrator Plus 5301).

### CellEKT workflow

The optimized chemical proteomics workflow incorporates the improvements made based on the chemical proteomics optimization experiments. Cells were treated with kinase inhibitor and probe as described in the section ‘Cell treatment for chemical proteomics’ and snap-frozen. 1.5×10^6^ HEK293T cells and 7×10^6^ MV4-11 cells were used to receive around 0.75 mg of final protein after cell lysis for each condition. Kinase inhibitors are tested in a full dose response manner at 6 different concentrations from 0.1 nM to 10,000 nM with each concentration in biological duplicates n = 2. Additionally, n = 3 – 4 biological replicates as peptide library samples are included by treating cells with 10 µM probe. All resulting cell pellets were lysed by vortexing in 400 µL ice-cold lysis buffer containing 50 mM HEPES pH 7.5, 150 mM NaCl, 1 mM MgCl_2_, 0.1% (w/v) Triton X-100, 1X cOmplete™ EDTA-free Protease Inhibitor Cocktail (Roche) and 25 U/mL benzonase (sc-202391, Santa Cruz). Protein concentrations were measured by BCA assay (Thermo Fisher), equalized and 360 µL lysate was transferred to room-temperature 2 mL Eppendorf tubes containing 40 µL 10% SDS (1% final), vortexed and left at room temperature.

The lysates were subjected to a click reaction for 1 h with freshly prepared click mix (31.2 µL per sample: 16 µL 25 mM CuSO_4_ in MilliQ, 3.2 µL 25 mM THPTA in DMSO, 4 µL 10 mM biotin-N_3_ in DMSO and 8 µL 50 mM TCEP.HCl freshly dissolved in DPBS). Proteins were precipitated by addition of HEPES/EDTA buffer (80 µL, 50 mM HEPES, 50 mM EDTA, pH 7.5), MeOH (666 µL), CHCl_3_ (166 µL) and MilliQ (150 µL), vortexing after each addition. After spinning down (1,500 *g*, 5 min) the upper and lower layer were aspirated and the protein pellet was resuspended in MeOH (600 µL) by sonication (Qsonica Q700 Microplate Sonicator, 2 × 10 s pulses, 10% amplitude).

The protein was spun down (20,000 *g*, 5 min) and the supernatant was discarded. The proteins were redissolved in 0.5% SDS in PBS (500 µL) containing 5 mM DTT by heating (65 °C, 15 min, 800 rpm shaking). Samples were allowed to reach room temperature and treated with IAA (15 µL, 0.5 M in MilliQ, 10 mM final) for 30 min at room temperature. Samples were spun down briefly (20,000 *g*, 2 min), inspected for undissolved material and the supernatant was transferred to new 1.5 mL Eppendorf tubes containing prewashed beads.

Beads were prepared by mixing 10 µL high-capacity streptavidin agarose (Pierce, 20 µL of a 50% slurry) with 10 µL control agarose (Pierce, 20 µL of a 50% slurry) and washing the mixture twice with PBS + 0.5% SDS and once with PBS by vortexing, spinning down (3,000 *g*, 2 min) and discarding the supernatant each step. The beads were suspended in 500 µL PBS containing 10 mM DTT before addition of the protein sample.

After the sample and beads were combined, the mixture was agitated by shaking (1,000 rpm, 2 h). The beads were then spun down (3,000 *g*, 2 min), supernatant was discarded and the beads were washed (3 x PBS + 0.5% SDS, 3 x PBS) by vortexing, spinning down (3,000 *g*, 2 min) and discarding the supernatant each step. The beads were resuspended in MilliQ, transferred to low-binding tubes (Sarstedt), spun down (3,000 *g*, 2 min) and the supernatant was aspirated. Then, beads were suspended in digestion buffer (200 µL, 100 mM Tris pH 7.8, 100 mM NaCl, 1 mM CaCl_2_, 2% (v/v) acetonitrile, sequencing grade trypsin (Promega, 0.5 µg)) and incubated while shaking overnight (16 h, 37 °C, 1,000 rpm). Trypsin was quenched by addition of a solution of 10% (v/v) aqueous FA, the beads were spun down (3,000 *g*, 2 min) and the peptides in the supernatant were desalted using C18 StageTips^48^ preconditioned with 50 µL MeOH, 50 µL of 0.5% (v/v) FA in 80% (v/v) acetonitrile/MilliQ (solution B) and 50 µL 0.5% (v/v) FA in MilliQ (solution A) by centrifugation (600 *g*, 1-2 min)). The peptides on the StageTips were washed with solution A (100 µL) and eluted into new low-binding tubes using solution B (100 µL). Samples were concentrated using an Eppendorf speedvac (Eppendorf Concentrator Plus 5301).

### Nano-LC-MS settings for pulldown samples

Desalted peptide samples were reconstituted in 30 µL LC-MS solution (97:3:0.1 H_2_O, ACN, FA) containing 10 fmol/µL yeast enolase digest (cat. 186002325, Waters) as injection control. Injection amount was titrated using a pooled quality control sample to prevent overloading the nanoLC system and the automatic gain control (AGC) of the QExactive mass spectrometer.

The desalted peptides were separated on a UltiMate 3000 RSLCnano system set in a trap-elute configuration with a nanoEase M/Z Symmetry C18 100 Å, 5 µm, 180 µm x 20 mm (Waters) trap column for peptide loading/retention and nanoEase M/Z HSS C18 T3 100 Å, 1.8 µm, 75 µm x 250 mm (Waters) analytical column for peptide separation. The column was kept at 40 °C in a column oven.

Samples were injected on the trap column at a flow rate of 15 µL/min for 2 min with 99% mobile phase A (0.1% FA in ULC-MS grade water (Biosolve)), 1% mobile phase B (0.1% FA in ULC-MS grade acetonitrile (Biosolve)) eluent. The 85 min LC method, using mobile phase A and mobile phase B controlled by a flow sensor at 0.3 µL/min with average pressure of 400-500 bar (5,500-7,000 psi), was programmed as gradient with linear increment to 1% B from 0 to 2 min, 5% B at 5 min, 22% B at 55 min, 40% B at 64 min, 90% B at 65 to 74 min and 1% B at 75 to 85 min. The eluent was introduced by electro-spray ionization (ESI) via the nanoESI source (Thermo) using stainless steel Nano-bore emitters (40 mm, OD 1/32”, ES542, Thermo Scientific).

The QExactive HF was operated in positive mode with data dependent acquisition without the use of lock mass, default charge of 2+ and external calibration with LTQ Velos ESI positive ion calibration solution (88323, Pierce, Thermo) every 5 days to less than 2 ppm. The tune file for the survey scan was set to scan range of 350 – 1,400 m/z, 120,000 resolution (m/z 200), 1 microscan, automatic gain control (AGC) of 3e6, max injection time of 100 ms, no sheath, aux or sweep gas, spray voltage ranging from 1.7 to 3.0 kV, capillary temp of 250 °C and an S-lens value of 80. For the 10 data dependent MS/MS events the loop count was set to 10 and the general settings were resolution to 15,000, AGC target 1e5, max IT time 50 ms, isolation window of 1.6 m/z, fixed first mass of 120 m/z and normalized collision energy (NCE) of 28 eV. For individual peaks the data dependent settings were 1.00e3 for the minimum AGC target yielding an intensity threshold of 2.0e4 that needs to be reached prior of triggering an MS/MS event. No apex trigger was used, unassigned, +1 and charges >+8 were excluded with peptide match mode preferred, isotope exclusion on and dynamic exclusion of 10 sec.

In between experiments, routine wash and control runs were done by injecting 5 µl LC-MS solution containing 5 µL of 10 fmol/µL BSA or enolase digest and 1 µL of 10 fmol/µL angiotensin III (Fluka, Thermo)/oxytocin (Merck) to check the performance of the platform on each component (nano-LC, the mass spectrometer (mass calibration/quality of ion selection and fragmentation) and the search engine).

### MaxQuant processing

Raw files were analyzed with MaxQuant (Version 1.6.17.0 for the cell line screen, version 2.0.1.0 for all other experiments). The following changes were made to the standard settings of MaxQuant: Label-free quantification was enabled with an LFQ minimal ratio count of 1. Match between runs and iBAQ quantification were enabled. Searches were performed against a Uniprot database of the human proteome (UPID: UP000005640, downloaded August 14^th^, 2020) including yeast enolase (P00924). The “peptides.txt” and “proteingroups.txt” files were used for further analysis in Microsoft Excel, GraphPad Prism 8.1.1 (GraphPad Software Inc) and KNIME Analytics Platform 4.5.2. ‘Probe-enriched kinases’ were defined by filtering the Uniprot ID for human reviewed proteins annotated with keyword “Kinase” [KW-0418] in UniProt (Table S1), a fold-change in LFQ of at least 2 between positive (XO44 treatment) and negative control (Vehicle treatment), 2 or more unique peptides detected and at most 1 missing LFQ value in the positive control replicates. For KNIME processing, enrichment was based on protein intensity instead of LFQ intensity as this afforded more relevant results.

### Data analysis in KNIME

The data analysis workflow was built in KNIME Analytics Platform 4.5.2 with two additional modules: KNIME Python Integration and the Vernalis KNIME nodes. The workflow is fully annotated to provide insight in the data analysis steps. A sample list template has been created in Microsoft Excel that provides a convenient user input for the automated workflow. The sample list template is used to cross-correlate .raw file names and treatment conditions, as well as define proteins of interest (POI). This sample list, together with the MaxQuant output, is read in by KNIME. Here, in brief, the MaxQuant output is sanitised by removal of reverse peptides and contaminants, and grouped identifications are split towards their first (predominant) identification. Proteins with fewer than 2 unique peptides (configurable) are removed. A Gaussian distribution is fitted through all log_2_ transformed LFQ values using Pythons SciPy norm.fit. This distribution is then transformed by shifting µ left 1.8σ and reducing σ by 70% as per default settings of Perseus developed by Jürgen Cox and colleagues^41^. This resulting normal distribution is used to impute missing values for enriched proteins. Enrichment of proteins in probe-treated positive controls vs. DMSO treated negative controls is calculated based on raw Intensity values, and all proteins with a ratio ≥ 2 are considered probe targets. Further quality filtering is performed by ensuring that the positive controls have at most 1 missing LFQ intensity value. For the resulting enriched proteins missing values are imputed, and LFQ intensities are normalised with respect to mean unimputed positive control values. Proteins where the highest concentration of inhibitor shows ≤ 60 % probe labeling are flagged for pIC_50_ fitting. The IC_50_ and Hill slope are fitted using Pythons SciPy optimize.curve_fit algorithm using the equation:

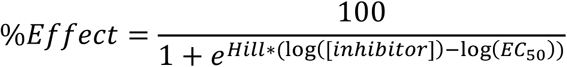

where *Hill* is the Hill-slope, [*inhibitor*] is the concentration of inhibitor used. This procedure and its output fits with error estimates is identical to standard IC_50_ curve fitting algorithms such as integrated in applications such as GraphPad Prism and relevant R-packages. Initial values for Hill slope and IC_50_ are provided as 1 and 1,000 nM, respectively. The Hill slope is constrained to be fit between 0 and 5. Determined pIC_50_s are classified as ‘Good’ if 0.3 ≤ Hill slope ≤ 3.0, σ(pIC_50_) ≤ 0.5 and pIC_50_ ≥ Max(p[inhibitor])-0.5 and as ‘Poor’ otherwise. If there are more than 6 imputed values for a dose response curve the pIC_50_ values are classified as ‘Poor’. The workflow outputs R^2^ values for each fit, which should be used only with caution as this metric is inappropriate for non-linear fits.^55^ It is only included to more closely match output by other statistical software. LFQ intensity histogram plots are generated highlighting each protein to visualise the relative abundance. MS/MS counts and unique peptide counts, averaged per replicate, are plotted per concentration of inhibitor to show available MS evidence for each plotted intensity. Finally, a phylogenetic tree is generated based on the seminal paper of Manning *et al.*^1^ and data visualization tools developed in the Phanstiel lab.^57^ All plots are exported as scaled vector graphic files, and together with the curve fits written to a Microsoft Excel file.

The data processing workflow together with installation instructions and template sample lists are provided on GitHub: https://github.com/APAJanssen/KNIME_MaxQuant_processing

### Cell treatment for chemical proteomics using a probe cocktail

One-third of the cells, specifically 0.5 × 10^6^ HEK293T cells and 4 × 10^6^ MV4-11 cells, were treated with XO44, ALX005, or ALX011 following the standard CellEKT workflow to assess the selectivity profile of dasatinib. At lysate level, the conditions were combined such that one-third of the lysate was treated with each probe per condition.

### Nano-LC-MS settings for probe cocktail samples

Fractionated peptide samples were analyzed using a nanoElute 2 LC system (Bruker) coupled to a timsTOF HT mass spectrometer (Bruker). 5 µL of sample was loaded on a trap column (PepMap C18, 5 mm x 0.3 mm, 5 µm, 100 Å, Thermo Scientific) followed by elution and separation on the analytical column (PepSep C18, 25 cm x 75 µm, 1.5 µm, 100 Å, Bruker). A gradient of 2 - 25% solvent B (0.1% FA in ACN) in 35 min, 25 - 32% B in 5 min, 32 - 95% in 5 min and 95% B for 10 min at a flow rate of 300 nL/min (all % values are v/v, water, TFA and ACN solvents were purchased from Biosolve, LC-MS grade). ZDV Sprayer 20 µm (Bruker) installed in the nano-electrospray source (CaptiveSpray source, Bruker) was used with following source parameters: 1,600 V of capillary voltage, 3.0 L/min of dry gas, and 180 °C of dry temperature. The MS data was acquired in DDA-PASEF mode with an ion mobility window of 0.85 to 1.35 Vs/cm^2^ in a mass range from 100 m/z to 1,700 m/z with charge states from 0 to 5+. The dual TIMS analyzer was utilized under a fixed duty cycle, incorporating a 100 ms ramp time, resulting in a total cycle time of 1.17 s. Precursors that reached a target intensity of 20,000 (intensity threshold 2,500) were selected for fragmentation and dynamically excluded for 0.4 min (exclusion window: mass width 0.015 m/z; 1/K0 width 0.015 Vs/cm^2^). The collision energy was set to 20 eV at 0.6 Vs/cm^2^ and 59 eV at 1.6 Vs/cm^2^. The 1/K0 values in between were interpolated linearly and kept constant above or below. The quadrupole isolation width was set to 2 m/z for 700 m/z and to 3 m/z for 800 m/z. Isolation width was constant except for linear interpolation between specified points. For calibration of the TIMS elution voltage, the Agilent ESI-Low Tuning Mix was used with three selected ions (m/z, 1/K0: 622.0290, 0.9915; 922.0098, 1.1986; 1221.9906, 1.3934). Mass calibration is performed with Na Formate in HPC mode.

### Data processing

The raw TIMS data (.d folders) were directly loaded into FragPipe (version 22.0). The FragPipe interface was used with MSFragger^58–60^ (version 4.1), IonQuant (version 1.10.27) and philosopher (version 5.1.1)^61^. The analysis was performed with the standard LFQ-MBR workflow. Searches were performed against a Uniprot database of the human proteome (UPID: UP000005640, downloaded August 14^th^, 2020) including yeast enolase (P00924). The “combined_protein.tsv” file was used for further analysis using our standard KNIME data analysis workflow with slight adjustments due to the different output. Instead of filtering for unique peptides, the output is filtered for unique spectra. Removal of reverse peptides and contaminants is no longer necessary. DMSO-treated negative vs positive control (probe treated) are calculated based on raw Intensity values, and all proteins with a ratio ≥ 10 are considered probe targets.

### Cell lysate processing for phosphoproteomics

20×10^6^ MV4-11 cells were treated with vehicle or midostaurin in 2-4 biological replicates and incubated at 37°C for 60 min. The treatment and preparation of cell pellets were performed as detailed in the previous section ’Cell treatment for chemical proteomics,’ with the exception that no probe was used and the probe incubation step was omitted

### Sample preparation for phosphoproteomics

For phosphotyrosine (pY) enrichment, a total of 5 mg of protein per sample was used. A lysate of HCT116 (colon carcinoma cell line) was taken along as workflow control. Protein samples were reduced with 5 mM DTT (dithiothreitol) for 30 minutes at 55 °C followed by alkylation using 10 mM IAA (iodoacetamide) for 15 min at room temperature. Lysates were then diluted using three volumes of 20 mM HEPES to reduce the concentration of urea to 2 M. Samples were digested at room temperature overnight using 1:100 (m/m) sequencing grade trypsin (cat. no. V5111, Promega, USA).

After digestion, TFA was added to reach 0.1% concentration, tubes were cooled to 0 °C and precipitate was pelleted by centrifugation (5,000 g, 5 min). The peptides in the supernatant were desalted using Oasis HLB columns (500 mg capacity, Waters), eluted in 0.1% TFA, 80% ACN and lyophilized. Lyophilized phosphopeptides were dissolved in IAP buffer (20 mM Tris-HCl pH 7.2, 10 mM sodium phosphate and 50 mM NaCl) and incubated with PTMScan pY antibody-conjugated beads (p-Tyr-1000, Cell Signaling Technology, USA) at a ratio of 4 µL slurry per mg protein at 4 °C for 2 h as previously described.^43^ Then, beads were pelleted by centrifugation (2,000 g, 30 s), supernatant was removed and beads were washed twice with cold IAP buffer and thrice with Milli-Q water. Phosphopeptides were eluted from the beads using two 50 µL volumes of 0.15% TFA and directly loaded on Evotips (Evosep, Denmark). Peptides were concentrated in a vacuum centrifuge at 45 °C, redissolved in 0.1% formic acid and an aliquot was loaded on Evotips.

### Data acquisition by LC-MS/MS

#### LC analysis

Peptides were separated via nanoflow reversed-phase liquid chromatography using standardized gradients on an Evosep One liquid chromatography system (Evosep, Denmark) with 0.1% FA and 0.1% FA/99.9% ACN as the mobile phases. The 30 samples per day (SPD) method was used in combination with a 15 cm x 150 μm reverse-phase column packed with 1.5 μm C18-beads (Bruker Daltonics) connected to a 20 μm ID fused silica emitter (Bruker Daltonics). Peptides were introduced to a timsTOF HT (Bruker Daltonics) using a nano-electrospray ion source (Captive spray source, Bruker Daltonics) with spray voltage set to 1500 V.

#### MS analysis

Peptides were analyzed on a TimsTOF HT running in DDA-PASEF mode. The ramp time was set to 100 ms and ten PASEF scans were acquired per topN acquisition cycle, resulting in a cycle time of 1.16s. Precursors with a mass range from 100 m/z to 1700 m/z, ion mobility range from 1.5 to 0.7 Vs cm−2¬¬, and charge states from 0 (unassigned) to 5+ were analyzed. The intensity threshold was set to 2,500 arbitrary units (a.u.) and target value to 20,000 a.u. Precursors that reached this target value or full scheduling capacity were excluded for 0.2 min. Singly charged precursors were filtered out based on their m/z-ion mobility position. Precursors with a mass below 700 Da were isolated with a quadrupole isolation width of 2 Th, precursors above 700 Da with a width of 3 Th. Collision energy was linearly decreased from 59 eV at 1.4 Vs cm-2 to 20 eV at 0.6 Vs cm-2. For all experiments the ion mobility dimension was calibrated linearly using three selected ions of the Agilent ESI LC/MS Tuning Mix [m/z, 1/K0: (322.0481, 0.7318 Vs cm−2), (622.0289, 0.9848 Vs cm−2), (922.0097, 1.1895 Vs cm−2)].

#### Phosphopeptide and -site identification and quantification by LC-MS/MS

The DDA-PASEF files of IMAC- and pTyr IP-enriched peptides searched separately against the Swissprot human FASTA file (downloaded March 2023, canonical and isoforms; 42,420 entries) using MaxQuant 2.3.1.0. Enzyme specificity was set to trypsin, and up to two missed cleavages were allowed. Cysteine carboxamidomethylation (+57.021464 Da) was treated as fixed modification and serine, threonine, and tyrosine phosphorylation (+79.966330 Da), methionine oxidation (+15.994915 Da), and N-terminal acetylation (+42.010565 Da) as variable modifications. Match between runs was enabled. For calculation of kinase INKA scores^43^, phosphopeptide MS/MS spectral counts were calculated from the MaxQuant evidence file using Rand used for substrate-centric inference of kinase activity on the basis of kinase–substrate relationships which are either experimentally observed (PhosphoSitePlus, “PSP”) or algorithm-predicted using sequence motif and protein–protein network information (NetworKIN, “NWK”). After literature analysis, NPM1 was excluded as kinase substrate. For data representation and analysis, biological replicates were averaged.

### Chemical Synthesis

#### Materials and General Methods

All reagents were purchased from chemical suppliers (Abovchem, Ambeed, BLD, Fisher Scientific, Fluorochem, Merck Sigma-Aldrich) and used without further purification. Solvents (Honeywell, VWR, Biosolve) indicated with “dry” were stored on activated 3 Å (EtOH, MeCN) or 4 Å (other solvents) molecular sieves (8 to 12 mesh, Acros Organics).

Microwave reactions were performed in a Biotage Initiator+ reactor. Reactions were monitored by thin layer chromatography (TLC, silica gel 60, UV_254_, Macherey-Nagel, ref: 818333) and compounds were visualized by UV absorption (254 nm and/or 366 nm) or spray reagent (permanganate (5 g/L KMnO_4_, 25 g/L K_2_CO_3_)) followed by heating. Alternatively, reactions were monitored by liquid chromatography-mass spectrometry (LCMS), either on a Thermo Finnigan (Thermo Finnigan LCQ Advantage MAX ion-trap mass spectrometer (ESI+) coupled to a Surveyor HPLC system (Thermo Finnigan) equipped with a Nucleodur C18 Gravity column (50×4.6 mm, 3 µm particle size, Macherey-Nagel)) or a Thermo Fleet (Thermo LCQ Fleet ion-trap mass spectrometer (ESI+) coupled to a Vanquish UHPLC system). LCMS eluent consisted of H_2_O in 0.1% TFA (aq.) and MeCN in 0.1% TFA (aq.). LCMS methods were as follows: 0.5 min cleaning with starting gradient, 8 min using specified gradient (linear), 2 min cleaning with 90% MeCN in 0.1% TFA (aq.). LCMS data is reported as follows: instrument (Finnigan or Fleet), gradient (% MeCN in 0.1% TFA (aq.)), retention time (t_r_) and mass (as m/z: [M+H]^+^). Purity of final compounds was determined to be ≥ 95% by integrating UV intensity of spectra generated by either of the LCMS instruments.

^1^H, ^13^C and ^19^F NMR spectra were recorded on a Bruker AV300 (300 (^1^H) and 75 MHz (^13^C), respectively), Bruker AV400 (400, 101 and 376 MHz, respectively), Bruker AV500 (500, 126 and 471 MHz, respectively) or Bruker AV600 (600 (^1^H) and 150 (^13^C), respectively) NMR spectrometer. NMR samples were prepared in deuterated chloroform (CDCl_3_), methanol (MeOD) or DMSO. Chemical shifts are given in ppm (δ) relative to residual protonated solvent signals (CDCl_3_ → δ 7.26 (^1^H), δ 77.16 (^13^C), MeOD → δ 3.31 (^1^H), δ 49.00 (^13^C), DMSO → δ 2.50 (^1^H), δ 39.52 (^13^C)). Data was processed by using MestReNova (v. 14) and is reported as follows: chemical shift (δ), multiplicity, coupling constant (*J* in Hz) and integration. Multiplicities are abbreviated as follows: s = singlet, br s = broad singlet, d = doublet, dd = doublet of doublets, t = triplet, dt = doublet of triplets, ddt = doublet of doublet of triplets, q = quartet, dq = doublet of quartets, m = multiplet. For some molecules about 1:1 rotamer and/or tautomer peaks were observed, resulting in additional peaks. For these compounds, chemical shifts were reported as ranges and multiplicity was denoted by “2x”, followed by the multiplicities specified above (i.e. 2x d = twice a doublet). The reported coupling constant corresponds to either of the multiplet peaks (of note, coupling constants were the same for both multiplet peaks).

Purification was done either by manual silica gel column chromatography (using 40-63 µm, 60 Å silica gel, Macherey-Nagel) or automated flash column chromatography on a Biotage Isolera machine (using pre-packed cartridges with 40-63 µm, 60 Å silica gel (4, 12, 25 or 40 g), Screening Devices).

High-performance liquid chromatography (HPLC) purifications were performed on either an Agilent 1200 preparative HPLC system (equipped with a Gemini C18 column (250×10 mm, 5 µm particle size, Phenomenex) coupled to a 6130 quadrupole mass spectrometer) or a Waters Acquity UPLC system (equipped with a Gemini C18 column (150×21 mm, 5 µm particle size, Phenomenex) coupled to a SQ mass spectrometer). Specified gradients for HPLC purifications (MeCN in 0.2% TFA (aq.)) were linear (5 mL/min for 12 min (Agilent) or 25 mL/min for 10 min (Waters)). High resolution mass spectrometry (HRMS) spectra were recorded through direct injection of a 1 µM sample either on a Thermo Scientific Q Exactive Orbitrap equipped with an electrospray ion source in positive mode coupled to an Ultimate 3000 system (source voltage = 3.5 kV, capillary temperature = 275 °C, resolution R = 240,000 at m/z 400, external lock, mass range m/z = 150-2,000).The eluent for HRMS measurements consisted of a 1:1 (v/v) mixture of MeCN in 0.1% formic acid (aq.) using a flow of 25 mL/min.

##### Scheme - Synthesis of pyrrolo[2,3-*d*]pyrimidine probes

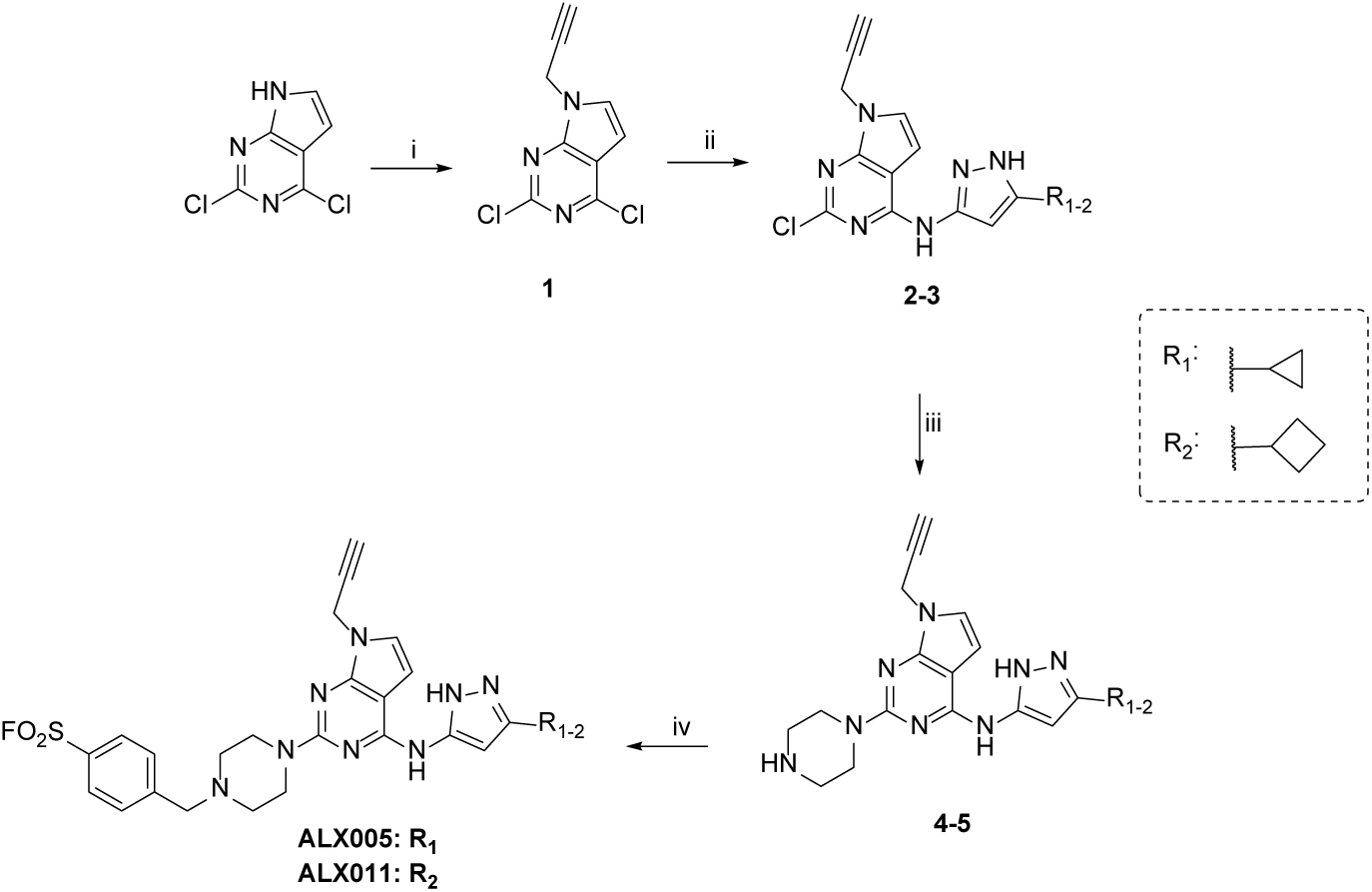

**Reagents and conditions:** i) K_2_CO_3_, 3-bromoprop-1-yne, DMF, RT, 96%. ii) 3-cyclopropyl-1*H*-pyrazol-5-amine (for **2**), 3-cyclobutyl-1*H*-pyrazol-5-amine (for **3**), DIPEA, THF, 60 – 120 °C, microwave irradiation, 17 - 58%. iii) piperazine, DMF, 100 – 130 °C, microwave irradiation, 26 - 29%. iv) 4-(bromomethyl)benzenesulfonyl fluoride, DIPEA, DMF, RT, 37 – 43%.

### Chemical Synthesis

#### General procedure A – Benzylation of piperazine derivatives 4-5

Piperazine analogue **4** or **5** (1 eq.), corresponding benzyl bromide (1.1 eq.) and DIPEA (4 eq.) were dissolved in DMF (molarity indicated) and the reaction mixture was stirred at RT for the indicated time. The reaction mixture was combined with sat. NaHCO_3_ (10 mL) and extracted with EtOAc (3×10 mL). The organic layer was washed with brine (10 mL), dried over MgSO_4_, filtered and concentrated. Purification was performed as indicated.

##### 4-((4-(4-((3-Cyclopropyl-1*H*-pyrazol-5-yl)amino)-7-(prop-2-yn-1-yl)-7*H*-pyrrolo[2,3-*d*]pyrimidin-2-yl)piperazin-1-yl)methyl)benzenesulfonyl fluoride (ALX005)

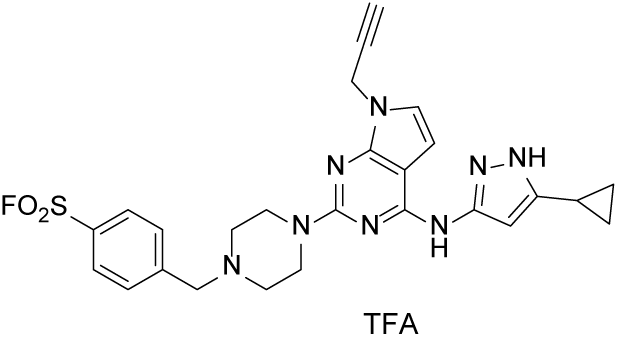

The title compound was synthesized from **4** (77.0 mg, 212 µmol) and 4-(bromomethyl)benzenesulfonyl fluoride (59.1 mg, 234 µmol) according to general procedure A (molarity: 0.08 M; reaction time: 1 h). The residue was purified by prep. HPLC to afford the product as a TFA salt (57.0 mg, 90 µmol, 43%). ^1^H NMR (400 MHz, MeOD) δ 8.19 (d, *J* = 8.5 Hz, 2H), 7.93 (d, *J* = 8.3 Hz, 2H), 7.25 (d, *J* = 3.8 Hz, 1H), 6.74 (d, *J* = 3.7 Hz, 1H), 5.94 (s, 1H), 4.96(d, *J* = 2.6 Hz, 2H), 4.54 (s, 2H), 4.15 – 4.08 (m, 4H), 3.46 (t, *J* = 5.2 Hz, 4H), 2.89 (t, *J* = 2.5 Hz, 1H), 2.05 – 1.92 (m, 1H), 1.13 – 1.06 (m, 2H), 0.81 (dt, *J* = 6.9, 4.7 Hz, 2H). (splitting up of carbons in aromatic region observed) ^13^C NMR (101 MHz, MeOD) δ 152.05, 151.73, 151.70, 150.77, 150.71, 149.14, 149.12, 148.93, 148.86, 139.14, 135.73, 135.48, 133.98, 130.28, 126.47, 101.40, 97.40, 92.29, 78.41, 74.96, 60.46, 52.28, 43.64, 34.40, 8.88, 7.61, 7.59. ^19^F NMR (471 MHz, DMSO) δ 65.92. LCMS (Fleet, 10% → 90%) t_R_ = 4.77 min; m/z: 535.42 [M+H]^+^. HRMS calculated for C_26_H_27_FN_8_O_2_S+H^+^: 535.20345, found 535.20300.

##### 4-((4-(4-((3-Cyclobutyl-1*H*-pyrazol-5-yl)amino)-7-(prop-2-yn-1-yl)-7*H*-pyrrolo[2,3-*d*]pyrimidin-2-yl)piperazin-1-yl)methyl)benzenesulfonyl fluoride (ALX011)

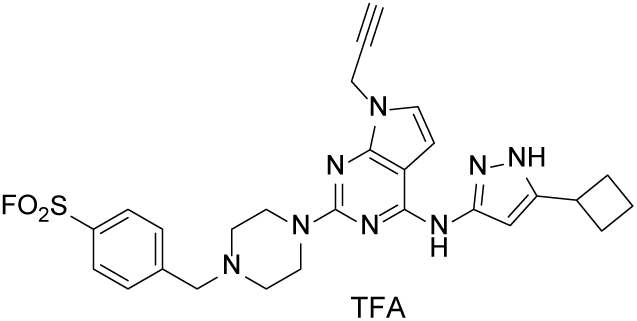

The title compound was synthesized from **5** (47.5 mg, 126 µmol) and 4-(bromomethyl)benzenesulfonyl fluoride (40.0 mg, 158 µmol) according to general procedure A (molarity: 0.08 M; reaction time: 1 h). The residue was purified by prep. HPLC to afford the product as a TFA salt (29.0 mg, 44.9 μmol, 37%). ^1^H NMR (400 MHz, MeOD) δ 8.18 (d, *J* = 8.4 Hz, 2H), 7.94 (d, *J* = 8.2 Hz, 2H), 7.26 (d, *J* = 3.8 Hz, 1H), 6.76 (d, *J* = 3.7 Hz, 1H), 6.12 (s, 1H), 4.97 (s, 2H), 4.52 (s, 2H), 4.19 – 4.02 (m, 4H), 3.63 (p, *J* = 8.6 Hz, 1H), 3.44 (t, *J* = 5.2 Hz, 4H), 2.90 (t, *J* = 2.5 Hz, 1H), 2.48 – 2.36 (m, 2H), 2.30 – 2.17 (m, 2H), 2.17 – 2.04 (m, 1H), 2.03 – 1.89 (m, 1H). ^13^C NMR (101 MHz, MeOD) δ 152.03, 151.78, 151.72, 151.58, 151.55, 149.07, 149.02, 139.50, 135.63, 135.37, 133.91, 130.25, 126.48, 101.46, 97.31, 93.36, 78.41, 74.96, 60.54, 52.31, 43.74, 34.41, 32.64, 30.01, 19.49. ^19^F NMR (471 MHz, DMSO) δ 65.93. LCMS (Fleet, 10% → 90%) t_R_ = 5.72 min; m/z: 549.13 [M+H]^+^. HRMS calculated for C_27_H_29_FN_8_O_2_S+H^+^: 549.21910, found 549.21871.

##### 2,4-Dichloro-7-(prop-2-yn-1-yl)-7*H*-pyrrolo[2,3-*d*]pyrimidine (1)

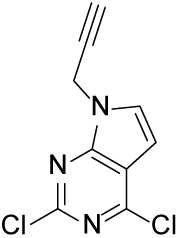

2,4-Dichloro-7*H*-pyrrolo[2,3-*d*]pyrimidine (1.00 g, 5.32 mmol) and K_2_CO_3_ (1.47 g, 10.6 mmol) were suspended in DMF (25.6 mL). 3-Bromoprop-1-yne (504 µL, 5.85 mmol) was added dropwise and the mixture was stirred at RT for 16 h. The reaction mixture was combined with H_2_O (50 mL) and extracted with EtOAc (3×50 mL). The organic layers were combined, washed with brine (3×50 mL), dried over MgSO_4_, filtered and concentrated to afford the product as an off-white solid (1.15 g, 5.09 mmol, 96%). ^1^H NMR (400 MHz, DMSO) δ 7.84 (d, *J* = 3.7 Hz, 1H), 6.75 (d, *J* = 3.6 Hz, 1H), 5.13 (d, *J* = 2.6 Hz, 2H), 3.52 (t, *J* = 2.5 Hz, 1H). ^13^C NMR (101 MHz, DMSO) δ 151.32, 151.19, 150.50, 131.76, 116.13, 99.75, 77.91, 76.40, 34.20. LCMS (Fleet, 10 → 90%): t_R_ = 7.42 min; m/z: 226.07 [M+H]^+^.

##### 2-Chloro-*N*-(3-cyclopropyl-1*H*-pyrazol-5-yl)-7-(prop-2-yn-1-yl)-7*H*-pyrrolo[2,3-*d*]pyrimidin-4-amine (2)

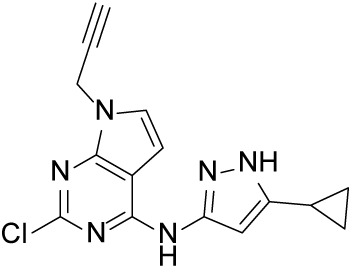

**1** (300 mg, 1.33 mmol), 3-cyclopropyl-1*H*-pyrazol-5-amine (254 mg, 2.06 mmol) and DIPEA (464 μL, 2.65 mmol) were dissolved in THF (2.13 mL) in a microwave vial and stirred at 120 °C for 12 h in the microwave. The reaction mixture was combined with H_2_O (10 mL) and extracted with DCM (3×10 mL). The organic layers were combined, dried over MgSO_4_, filtered and concentrated. The residue was purified by automated column chromatography (0.5% → 9% MeOH/DCM) to afford the product as a white solid (241 mg, 0.771 mmol, 58%). ^1^H NMR (400 MHz, DMSO) δ 12.18 (s, 1H), 10.33 (s, 1H), 7.28 (d, *J* = 3.6 Hz, 1H), 6.84 (br s, 1H), 6.41 (s, 1H), 4.97 (d, *J* = 2.6 Hz, 2H), 3.42 (t, *J* = 2.5 Hz, 1H), 1.98 – 1.87 (m, 1H), 0.97 – 0.90 (m, 2H), 0.75 – 0.67 (m, 2H). ^13^C NMR (101 MHz, DMSO) δ 153.89, 152.37, 150.00, 147.40, 145.47, 124.74, 101.88, 100.22, 94.47, 78.86, 75.63, 33.37, 7.70, 6.94. LCMS (Fleet, 10% → 90%) t_R_ = 5.60 min; m/z: 313.13 [M+H]^+^.

##### *N*-(3-Cyclobutyl-1*H*-pyrazol-5-yl)-2-(piperazin-1-yl)-7-(prop-2-yn-1-yl)-7*H*-pyrrolo[2,3-*d*]pyrimidin-4-amine (3)

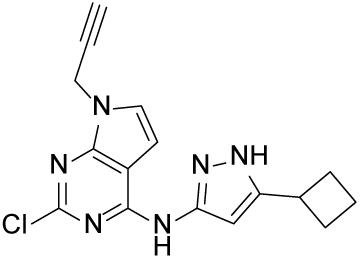

**1** (650 mg, 2.88 mmol), 3-cyclobutyl-1*H*-pyrazol-5-amine (592 mg, 4.31 mmol) and Et_3_N (802 μL, 5.75 mmol) were dissolved in MeOH (10.0 mL) in a microwave vial and stirred at 60 °C for 36 h in the microwave. The reaction mixture was combined with H_2_O (10 mL) and extracted with DCM (3×10 mL). The organic layers were combined, dried over MgSO_4_, filtered and concentrated. The residue was purified by automated column chromatography (0.5% → 9% MeOH/DCM) to afford the product as a white solid (162 mg, 0.496 mmol, 17%). ^1^H NMR (400 MHz, DMSO) δ 12.24 (s, 1H), 10.36 (s, 1H), 7.29 (d, *J* = 3.6 Hz, 1H), 6.86 (br s, 1H), 6.55 (s, 1H), 4.97 (d, *J* = 2.6 Hz, 2H), 3.52 (p, *J* = 8.7 Hz, 1H), 3.42 (t, *J* = 2.5 Hz, 1H), 2.36 – 2.23 (m, 2H), 2.23 – 2.08 (m, 2H), 2.03 – 1.78 (m, 2H). ^13^C NMR (101 MHz, DMSO) δ 153.86, 152.40, 149.99, 147.39, 146.93, 124.72, 101.90, 100.24, 94.97, 78.87, 75.63, 3f3.38, 31.26, 29.04, 18.15. LCMS (Fleet, 10% → 90%) t_R_ = 6.06 min; m/z: 327.13 [M+H]^+^.

##### *N*-(3-Cyclopropyl-1*H*-pyrazol-5-yl)-2-(piperazin-1-yl)-7-(prop-2-yn-1-yl)-7*H*-pyrrolo[2,3-*d*]pyrimidin-4-amine (4)

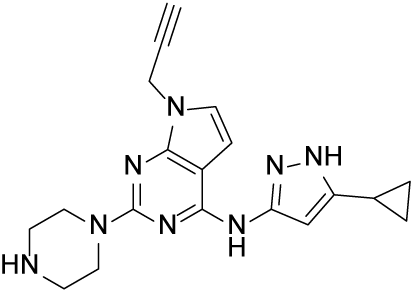

**2** (262 mg, 0.838 mmol) and piperazine (289 mg, 3.35 mmol) were dissolved in DMF (13.1 mL) in a microwave vial and the reaction mixture was stirred in the microwave at 100 °C for 18 h. The reaction mixture was combined with EtOAc (30 mL) and washed with sat. NH_4_Cl (30 mL), sat. NaHCO_3_ (30 mL), and brine (30 mL). The organic layer was dried over MgSO_4_, filtered and concentrated. The residue was purified by automated column chromatography (10% → 20% MeOH/DCM) to afford the product as yellow crystals (87 mg, 0.24 mmol, 29%). (NH of piperazine not observed) ^1^H NMR (400 MHz, DMSO) δ 12.00 (br s, 1H), 9.60 (s, 1H), 6.87 (d, *J* = 3.6 Hz, 1H), 6.66 (s, 1H), 6.32 (s, 1H), 4.83 (d, *J* = 2.5 Hz, 2H), 3.67 – 3.60 (m, 4H), 3.33 (t, *J* = 2.5 Hz, 1H), 2.79 – 2.72 (m, 4H), 1.94 – 1.83 (m, 1H), 0.98 – 0.89 (m, 2H), 0.70 – 0.62 (m, 2H). ^13^C NMR (101 MHz, DMSO) δ 158.70, 153.16, 151.72, 147.59, 145.87, 120.46, 99.89, 96.36, 93.19, 79.66, 74.86, 45.28, 45.03, 32.34, 7.89, 7.13. LCMS (Fleet, 10% → 90%) t_R_ = 4.25 min; m/z: 363.27 [M+H]^+^.

##### *N*-(3-Cyclobutyl-1*H*-pyrazol-5-yl)-2-(piperazin-1-yl)-7-(prop-2-yn-1-yl)-7*H*-pyrrolo[2,3-*d*]pyrimidin-4-amine (5)

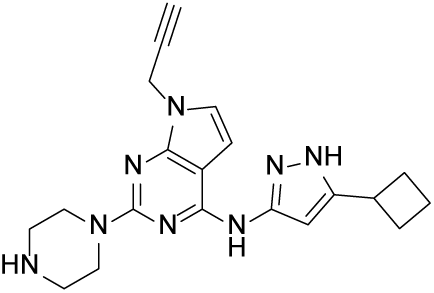

**3** (162 mg, 0.496 mmol) and piperazine (171 mg, 1.78 mmol) were dissolved in DMF (5.00 mL) in a microwave vial and the reaction mixture was stirred in the microwave at 130 °C for 18 h. The reaction mixture was combined with EtOAc (10 mL) and washed with sat. NH_4_Cl (10 mL), sat. NaHCO_3_ (10 mL), and brine (10 mL). The organic layer was dried over MgSO_4_, filtered and concentrated. The residue was purified by automated column chromatography (10% → 20% MeOH/DCM) to afford the product as a dark green oil (47.5 mg, 0.496 mmol, 26%). (NH of piperazine and NH of pyrazole not observed) ^1^H NMR (400 MHz, DMSO) δ 9.68 (s, 1H), 6.89 (d, *J* = 3.6 Hz, 1H), 6.69 (d, *J* = 3.7 Hz, 1H), 6.51 (s, 1H), 4.84 (d, *J* = 2.6 Hz, 2H), 3.78 – 3.71 (m, 4H), 3.49 (p, *J* = 8.5 Hz, 1H), 3.34 (t, *J* = 2.5 Hz, 1H), 2.90 – 2.83 (m, 4H), 2.36 – 2.23 (m, 2H), 2.19 – 2.05 (m, 2H), 2.02 – 1.80 (m, 2H). 13C NMR (101 MHz, DMSO) δ 158.61, 153.26, 151.70, 147.71, 147.52, 120.57, 99.97, 96.52, 94.36, 79.66, 74.89, 44.78, 44.43, 32.39, 31.40, 29.10, 18.26. LCMS (Fleet, 10% → 90%) t_R_ = 3.82 min; m/z: 377.17 [M+H]^+^.

### SIK2 NanoBRET™ Target Engagement (TE) Assay

Monoclonal HEK293 stale cell lines, expressing NanoLuc-tagged human/mouse SIK2 were established by transfection with corresponding plasmids, followed by selection with 700 ug/mL G418. The NanoBRET™ TE Assays were performed with commercial K4 (Promega) or in-house synthesized tracers according to the manufacture’s protocol (NanoBRET™ TE Intracellular Kinase Assay Kit, Catalogue # N2521, Promega).

### SIK2 biochemical assay with RapidFire-MS readout

In the presence of SIK2 and ATP the CHK-peptide (KKKVSRSGLYRSPSMPENLNRPR with C-terminal arginine amide modification) was phosphorylated at one of the four feasible serine residues. Only one phosphorylation is observed under the assay conditions. 60 nL of each compound dilution series (12 or 24 point; dilution factor 3, generally 30 μΜ to 170 pM) in DMSO were transferred by acoustic dispensing to the assay plate and 30 min pre-incubated (ambient temperature) after the addition of 5 μL SIK2 (0.5 nM f.c.) in assay buffer (12.5 mM HEPES (pH 7.0), 10 mM magnesium acetate, 0.005 % BSA). 5 µL substrate solution (10 μΜ f.c. CHK-peptide, 100 μΜ ATP f.c.) for SIK2 in assay-buffer were added and incubated at ambient temperature for 45 min. 40 μL 0.125% formic acid in water was added to quench the reaction. RapidFire (RF) Mass Spectrometry was utilized for data generation as described below. The multiple charged species (3-5 charges) for the phosphorylated and non-phosphorylated form measured by MRM (Multiple Reaction Monitoring; API5000 or 6500+) or EIC (Extracted Ion Current; QToF) were summed up and the ratio calculated (sum phosphorylated species / sum all species) for data evaluation. Normalization was performed by Genedata software based on the non-inhibition control DMSO and the commercially available SIK inhibitor @ 1 μΜ YKL-05-099 (CAS number 1936529-65-5). The results of the assay are expressed in half-maximal inhibitory concentrations (IC50s).

RapidFire setup: Samples were aspirated by vacuum for max. 600 ms and loaded to C4-cartridge (Agilent; #G9203A) for 3000ms at 1.5 mL/min with 0.1% formic acid in water. Afterwards samples were transferred to the API5000 (API6500+) or QToF mass spectrometer for 4,000 ms at 1.25 mL/min 5 with 90% acetonitrile; 10% water; 0.007% TFA; 0.093 formic acid. The cartridge was reconditioned for additional 500 ms with 0.1% formic acid in water.

MS-Setup Sciex API5000/API6500+: All MS analyses using the following MS-setup in MRM mode: Electrospray positive; Ion Spray Voltage: 4000V; Temperature: 550 °C; Collision Gas: 5; Curtain Gas: 15; Gasl: 40; Gas 2: 42; io EP: 10

### Optimization of lysis and CuAAC conditions

To investigate post-lysis labeling, cell pellets of THP1 cells treated with staurosporine (1 µM, 30 min) and XO44 (1 µM, 30 min) were lysed completely in ice-cold lysis buffer containing 50 mM HEPES pH 7.5, 150 mM NaCl, 1 mM MgCl_2_, 0.1% (w/v) Triton X-100, 1X cOmplete™ EDTA-free Protease Inhibitor Cocktail (Roche) and 25 U/mL benzonase (sc-202391, Santa Cruz). The lysate was aliquoted in several Eppendorf tubes and treated with 0/0.1/1% SDS as well as additional staurosporine or vehicle (0.1% DMSO). Lysates were kept on ice (0 °C) or warmed to 37 °C for 2.5 h. SDS concentrations were equalized, all lysates were allowed to reach room temperature and lysates were subjected to a click reaction for 30 min with freshly prepared click mix (8.26 µL per 80 µL sample: 4.38 µL 25 mM CuSO_4_ in MilliQ, 0.88 µL 25 mM THPTA in DMSO, 0.2 µL 2 mM AF647-N_3_ (Thermo Fisher) in DMSO and 2.6 µL 250 mM sodium ascorbate in MilliQ). Samples were quenched with 4X Laemmli buffer and resolved by SDS-PAGE (10% acrylamide gel, ±80 min, 180 V) along with protein marker (PageRuler™ Plus, Thermo Fisher). In-gel fluorescence was measured in the Cy3- and Cy5-channel (Chemidoc™ MP, Bio-Rad) and the gels were subsequently stained with Coomassie and imaged as a loading control for normalization of fluorescence intensity.

To investigate the effect of lysis buffer components on click chemistry, cell pellets of THP1 cells treated with FP-alkyne (10 µM, 30 min) or vehicle were lysed completely in ice-cold lysis buffer A (250 mM sucrose, 20 mM HEPES pH 7.5, 1 mM MgCl_2_, 1X cOmplete™ EDTA-free Protease Inhibitor Cocktail (Roche) and 25 U/mL benzonase (sc-202391, Santa Cruz) or B (0.1% (w/v) Triton X-100, 20 or 50 mM HEPES pH 7.5, 150 mM NaCl, 1 mM MgCl_2_, 1X cOmplete™ EDTA-free Protease Inhibitor Cocktail (Roche) and 25 U/mL benzonase (sc-202391, Santa Cruz). The lysate was aliquoted in several Eppendorf tubes and SDS was added to the indicated concentration. Two samples were incubated at 37 °C for 1 h before addition of SDS to investigate post-lysis labeling. Lysates were treated with freshly prepared click mix containing either NaAsc or TCEP as reducing agent and different ratios of TCEP to CuSO_4_. Generally, per 40 µL of lysate was added click mix containing 2.19 µL 25 mM CuSO_4_ in MilliQ, 1.3 µL 250 mM sodium ascorbate in MilliQ or 1.1 µL 50 mM TCEP.HCl freshly dissolved in DPBS, 0.44 µL 25/100/200 mM THPTA in DMSO and 0.44 µL Cy5-N_3_. Samples were incubated for 1 h at room temperature before quenching with 4X Laemmli buffer, samples were resolved on SDS-PAGE gel and in-gel fluorescence was analyzed as described previously.

To investigate the compatibility of high SDS concentrations with TAMRA-biotin-N_3_ (Click Chemistry Tools), cell pellets of MV4-11 cell treated with XO44 (1 µM, 30 min) were lysed completely in ice-cold lysis buffer containing 50 mM HEPES pH 7.5, 150 mM NaCl, 1 mM MgCl_2_, 0.1% (w/v) Triton X-100, 1X cOmplete™ EDTA-free Protease Inhibitor Cocktail (Roche) and 25 U/mL benzonase (sc-202391, Santa Cruz). The lysate was aliquoted in several Eppendorf tubes and treated with 0.2 or 1% SDS at rt for 1 h, after which freshly prepared click mix was added (3.12 µL per 40 µL lysate: 1.6 µL 25 mM CuSO_4_ in MilliQ, 0.32 µL 25 mM THPTA in DMSO, 0.4 µL 2.5 or 10 mM TAMRA-biotin-N_3_ in DMSO and 0.8 µL 50 mM TCEP.HCl freshly dissolved in DPBS). Samples were incubated for 30 min or 1 h at room temperature before quenching with 4X Laemmli buffer, samples were resolved on SDS-PAGE gel and in-gel fluorescence was analyzed as described previously.

### Optimization of pull-down conditions

Cell pellets of MV4-11 cell treated with XO44 (1 µM, 30 min) were lysed completely in ice-cold lysis buffer containing 50 mM HEPES pH 7.5, 150 mM NaCl, 1 mM MgCl_2_, 0.1% (w/v) Triton X-100, 1X cOmplete™ EDTA-free Protease Inhibitor Cocktail (Roche) and 25 U/mL benzonase (sc-202391, Santa Cruz). Protein concentration was determined using BCA assay (Thermo Fisher) and set to 1.5 mg/mL. 400 µL aliquots lysate containing 0.2% SDS were made, allowed to reach room temperature and subjected to a click reaction for 30 min with freshly prepared click mix (7.8 µL per 100 µL lysate: 4 µL 25 mM CuSO_4_ in MilliQ, 0.8 µL 25 mM THPTA in DMSO, 1 µL 10 mM TAMRA-biotin-N_3_ in DMSO and 2 µL 50 mM TCEP/HCl freshly dissolved in DPBS). Proteins were precipitated by addition of HEPES/EDTA buffer (80 µL, 50 mM HEPES, 50 mM EDTA, pH 7.5), MeOH (666 µL), CHCl_3_ (166 µL) and MilliQ (150 µL), vortexing after each addition. After spinning down (1,500 g, 10 min) the upper and lower layer were aspirated and the protein pellet was resuspended in MeOH (600 µL) by sonication (Qsonica Q700 Microplate Sonicator, 2 × 10 s pulses, 10% amplitude). The proteins were spun down (20,000 *g*, 5 min) and the supernatant was discarded. The proteins were dissolved in 200 µL PBS containing 0.5% SDS and 20 mM DTT by heating to 65 °C for 15 minutes, after which the dissolved proteins were diluted with 200 µL PBS giving 1.5 mg/mL protein and 0.25% SDS (‘Input’ fraction). Avidin agarose (Pierce) or high-capacity Streptavidin agarose (Pierce) were prewashed twice with PBS + 0.5% SDS and once with PBS, divided over 1.5 mL tubes in indicated amounts, spun down (3,000 *g*, 2 min) and all liquid was aspirated without losing beads by using a gel-loading tip pushed to the bottom of the tube. The XO44-treated sample was divided over 100 µL aliquots (1/4^th^ of a regular pulldown sample). The sample was incubated with the beads under vigorous shaking (1,300 rpm) ensuring the beads were in suspension. After indicated time points, the beads were spun down (3,000 *g*, 2 min) and the supernatant was collected, denatured using Laemmli buffer and loaded on SDS-PAGE gel (‘Supernatant’ fraction). The beads were washed twice with PBS + 0.5% SDS and once with PBS, spun down (3,000 *g*, 2 min) and all liquid was aspirated without losing beads by means of a gel-loading tip pushed to the bottom of the tube. Proteins were eluted from the beads by boiling (5 min, 95 °C) in 2X Laemmli buffer containing 2 mM biotin (Pierce), beads were spun down (3000 *g*, 2 min) and sample was loaded on SDS-PAGE gel (‘Elution’ fraction).

### Determining target engagement by pull-down, elution and western blot

MV4-11 cells in log phase were spun down (300 *g*, 5 min) and resuspended at 1.0×10^6^ cells per mL in IMDM supplemented with 0.1% delipidated BSA (Merck) and 10 mM HEPES. The cells were treated with kinase inhibitor or vehicle (0.1% DMSO) for 1 h at 37 °C after which 1 or 10 µL XO44 or vehicle (0.1% DMSO) was added and the cells were incubated for another 25 min. The cells were spun down (300 *g*, 5 min, 37 °C), resuspended in ice-cold DBPS (1 mL), transferred to 1.5 mL Eppendorf tubes and spun down again (1,000 *g*, 5 min, 4 °C). The supernatant was aspirated and the cells were snap-frozen and stored at - 80 °C. Cell pellets were lysed completely in 400 µL ice-cold lysis buffer containing 50 mM HEPES pH 7.5, 150 mM NaCl, 1 mM MgCl_2_, 0.1% (w/v) Triton X-100, 1X cOmplete™ EDTA-free Protease Inhibitor Cocktail (Roche) and 25 U/mL benzonase (sc-202391, Santa Cruz). Protein concentrations were equalized and 360 µL lysate was transferred to 2 mL Eppendorf tubes containing 40 µL 10% SDS (1% final), vortexed and left at room temperature. The lysates were subjected to a click reaction for 1 h with freshly prepared click mix (31.2 µL per sample: 16 µL 25 mM CuSO_4_ in MilliQ, 3.2 µL 25 mM THPTA in DMSO, 4 µL 10 mM biotin-N_3_ in DMSO and 8 µL 50 mM TCEP/HCl freshly dissolved in DPBS). Proteins were precipitated by addition of HEPES/EDTA buffer (80 µL, 50 mM HEPES, 50 mM EDTA, pH 7.5), MeOH (666 µL), CHCl_3_ (166 µL) and MilliQ (150 µL), vortexing after each addition. After spinning down (1,500 *g*, 10 min) the upper and lower layer were aspirated and the protein pellet was resuspended in MeOH (600 µL) by sonication (Qsonica Q700 Microplate Sonicator, 2 × 10 s pulses, 10% amplitude). The proteins were spun down (20,000 *g*, 5 min) and the supernatant was discarded. The proteins were redissolved in PBS containing 0.5% SDS (200 µL) and 5 mM DTT (65 °C, 15 min, 1,000 rpm shaking). The samples were allowed to reach rt and transferred to new 1.5 mL Eppendorf tubes containing 30 µL prewashed avidin agarose slurry in 200 µL PBS. Samples were incubated under shaking for 2 h (800 rpm), spun down (3,000 *g*, 2 min) and the supernatant was discarded. Samples were washed (3 x PBS + 0.5% SDS, 1 x PBS) and all PBS was aspirated without losing beads by means of a gel-loading tip pushed to the bottom of the tube. Proteins were eluted from the beads by boiling (5 min, 95 °C) in 2X Laemmli buffer containing 2 mM biotin (Pierce), beads were spun down (3,000 *g*, 2 min) and sample was loaded on a 10% SDS-PAGE gel. Proteins were resolved (±80 min, 180 V) along with protein marker (PageRuler™ Plus, Thermo Fisher) and transferred to a 0.2 µm polyvinylidene difluoride membrane by Trans-Blot Turbo™ Transfer system (Bio-Rad). Membranes were washed with TBS (50 mM Tris pH 7.5, 150 mM NaCl) and blocked with 5% (w/v) milk in TBS-T (50 mM Tris pH 7.5, 150 mM NaCl, 0.05% (w/v) Tween-20) for 1 h at room temperature. Membranes were incubated with primary antibody rabbit-anti-FES (85704, Cell Signaling Technologies, 1:1000, o/n, 4 °C) in 5% (w/v) BSA in TBS-T. The membranes were then washed three times with TBS-T (5 min) and incubated with secondary goat anti-rabbit-HRP (sc-2030, Santa Cruz, 1:4,000 in 5% (w/v) milk in TBS-T, 3 h, rt) and washed three times with TBS-T and once with TBS. Membranes were developed in Clarity Western ECL Substrate (Bio-Rad) and chemiluminescence was detected on ChemiDoc™ MP (Bio-Rad) in the chemiluminescence channel and colorimetric channel for the protein marker. Images were processed using Image Lab 6.0.1 (BioRad).

## QUANTIFICATION AND STATISTICAL ANALYSIS

Quantification of protein abundance was done using the MaxLFQ algorithm for label-free quantitation (LFQ) in MaxQuant. Further normalization and fitting of inhibitor dose-response values in the KNIME workflow was done using SciPy using the Least squares method with trf optimizer. Details of replicates and data analysis for specific experiments can be found in the figure legends or in the methods section. The reported mean is equivalent to the average of the values determined for each of the replicates and the reported standard deviation is a measure of the variance relative to the determined mean.

