## Supplementary figures for "CellEKT: A robust chemical proteomics workflow to profile cellular target engagement of kinase inhibitors"

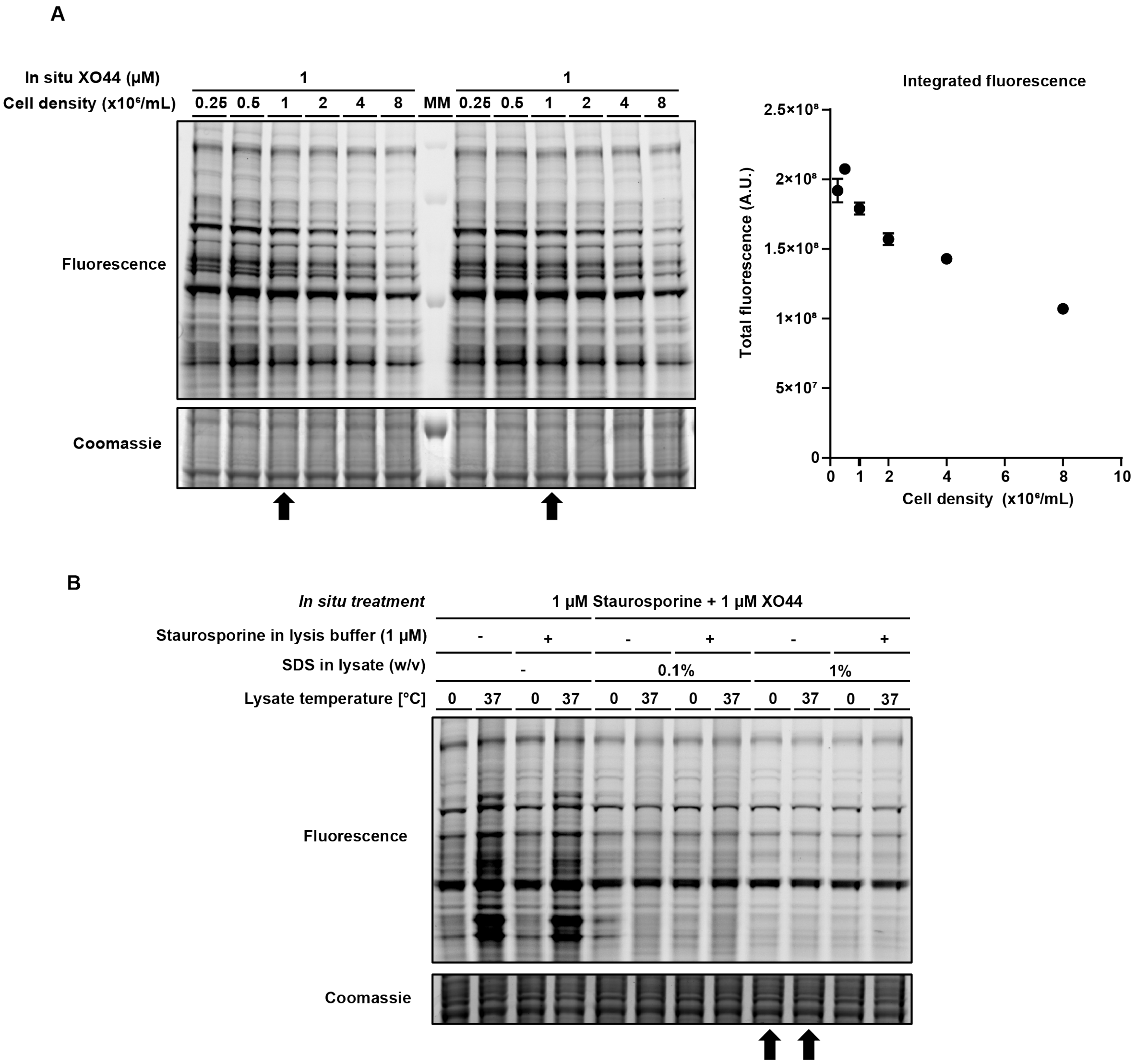


**Figure S1. Optimization of cell density and lysis conditions; Step 1 and 2. (A)** Treating cells at densities higher than 1x10$^6$/mL results in depletion of the probe. THP-1 cells at indicated density were treated with XO44 (1 µM, 30 min), harvested and lysed. XO44-labeled proteins were conjugated to AF647-$N_3$ using CuAAC chemistry and analyzed by SDS-PAGE and in-gel fluorescence scanning. Coomassie served as a protein loading control. **(B)** XO44 shows significant post-lysis labeling, which cannot be prevented with excess competitor in lysis buffer but can be prevented by protein denaturation. THP-1 cells were treated with staurosporine (1 µM, 30 min) and XO44 (1 µM, 30 min), harvested and lysed. Lysate was treated with 0/0.1/1% SDS and additional staurosporine (1 µM) or vehicle, and was either kept on ice or warmed to 37 °C for 2.5 h. XO44-labeled proteins were conjugated to AF647-$N_3$ using CuAAC chemistry and analyzed by SDS-PAGE and in-gel fluorescence scanning. Coomassie served as a protein loading control. Arrows indicate conditions which were deemed optimal.

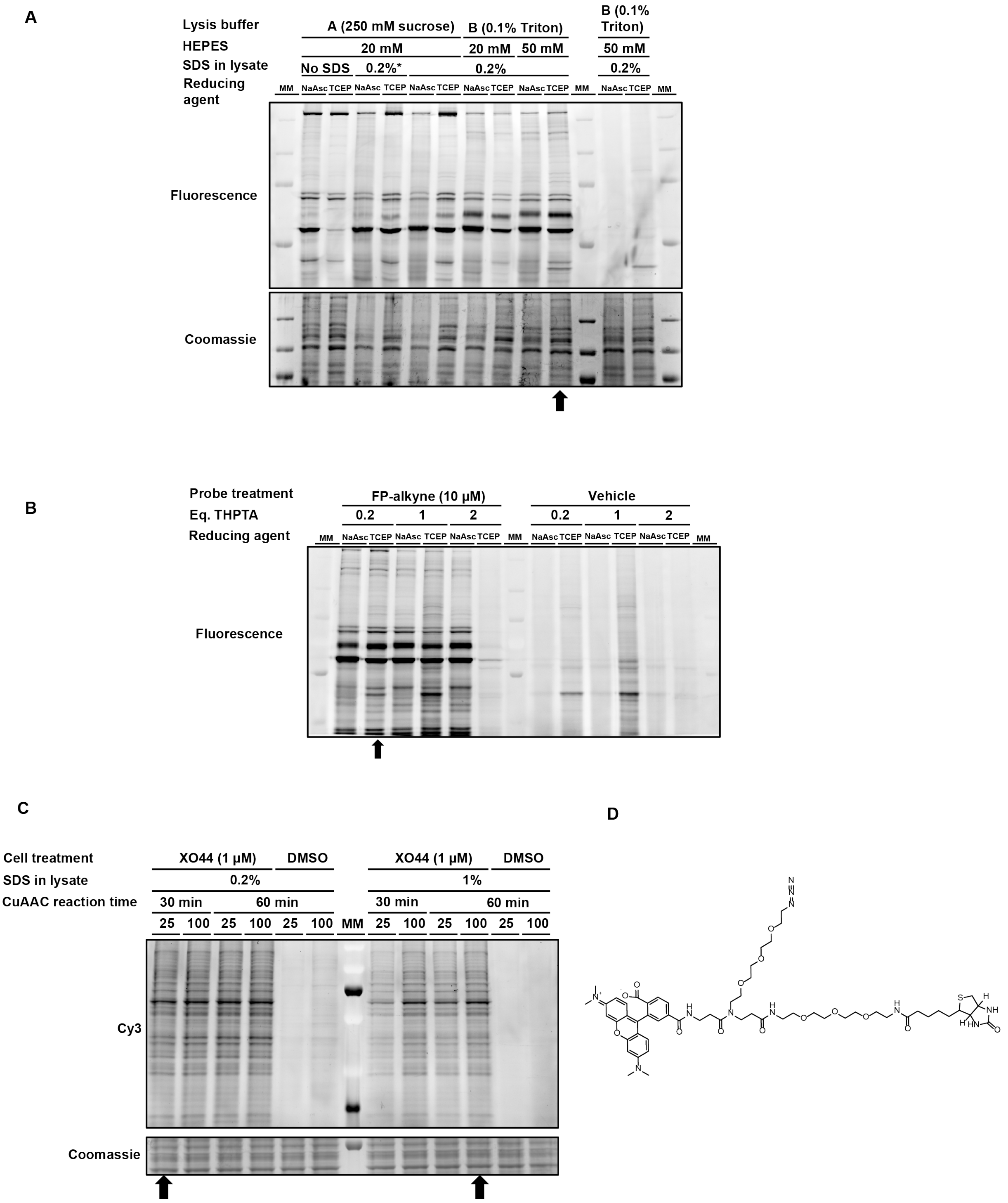


**Figure S2. CuAAC ligation was optimized using TCEP instead of sodium ascorbate and raising SDS concentration from 0.2% to 1% results in incomplete CuAAC labeling, which can be rescued by increasing azide reagent concentration or reaction time; Step 3.** THP-1 cells were treated with FP-alkyne (10 µM, 30 min), harvested and lysed in **(A)** indicated lysis buffer or **(B)** 0.2% SDS, 0.1% Triton X-100, 50 mM HEPES pH 7.5. Then, probe-labeled proteins were conjugated to Cy5-$N_3$ using CuAAC chemistry with indicated reducing agent and analyzed by SDS-PAGE and in-gel fluorescence scanning. Coomassie served as a protein loading control and demonstrates CuAAC-induced protein damage. *Lysate was incubated at 37 °C for 1 h before addition of SDS to investigate post-lysis labeling, which was not observed for FP-alkyne. Arrows indicate conditions which were deemed optimal. (**C**) Cell pellets of XO44-treated (1 µM, 30 min) MV4-11 cells were lysed, treated with 0.2 or 1% SDS and probe-labeled proteins were conjugated to TAMRA-biotin-N3 under indicated conditions. Proteins were analyzed by SDS-PAGE and in-gel fluorescence scanning. Coomassie served as a protein loading control. Arrows indicate CuAAC conditions which were deemed optimal for 0.2 and 1% SDS respectively. (**D**) Structure of commercially available TAMRA-biotin-$N_3$.

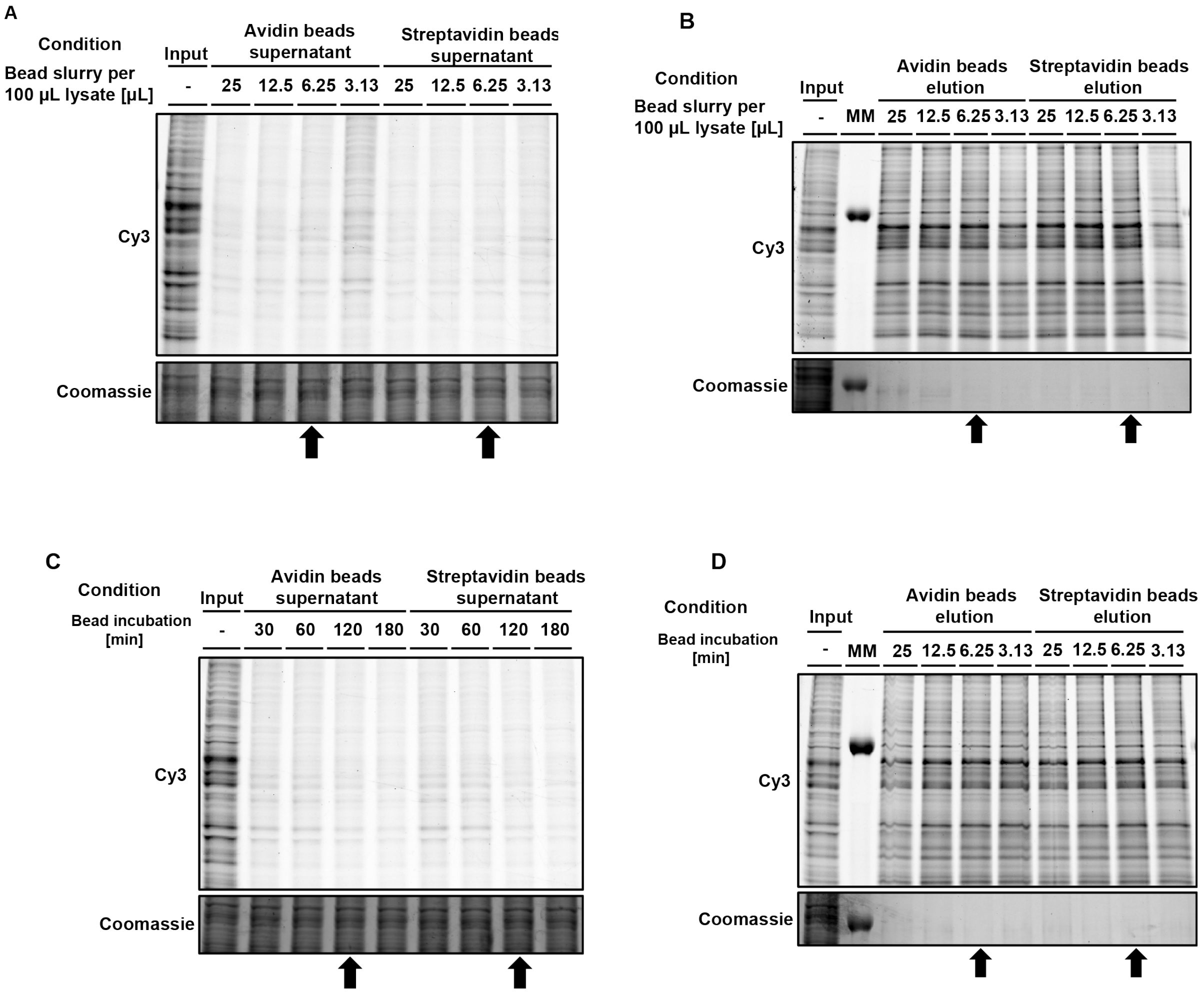


**Figure S3. Determination of optimal pulldown conditions of TAMRA-biotin-$N_3$ conjugated XO44-labeled proteins. For 100 µL lysate, 6.25 µL of either avidin or streptavidin agarose slurry is sufficient, lowering the volume further results in loss of beads during washing steps. Conjugation of biotinylated proteins is complete after 2 h of incubation; Step 4.** Lysate of XO44-treated MV4-11 cells was conjugated to TAMRA-biotin-$N_3$, proteins were precipitated using Bligh/Dyer and redissolved in 100 µL PBS containing 0.5% SDS (input fraction). Biotin-conjugated proteins were then pulled down by incubation with (strept)avidin agarose. After incubation, the bead supernatant was sampled (supernatant fraction), the beads were washed and labeled proteins were eluted using Laemmli buffer containing 2 mM biotin (bead elution fraction). Either **(A, B)** the amount and type of beads was varied and incubation time was kept constant (3 h), or **(C, D)** incubation time was varied and bead amount was kept constant (25 µL slurry). Arrows indicate conditions which were deemed optimal.

A

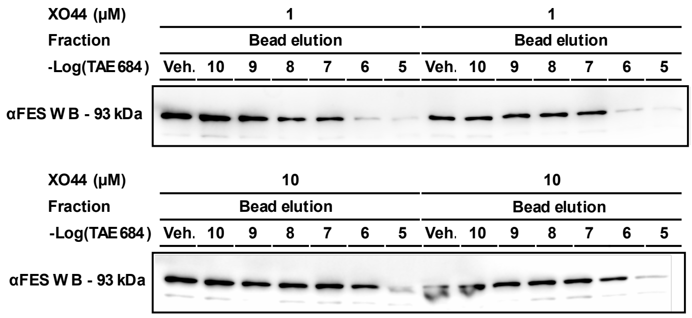


B

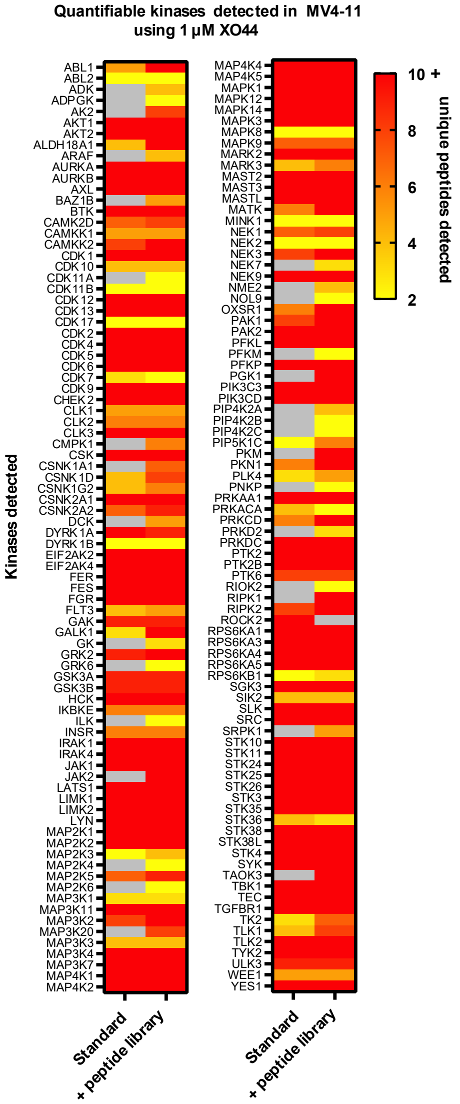


**Figure S4: Increasing the probe concentration leads to an underestimation of inhibitor target engagement. However, samples with high probe doses can be used to enhance kinome coverage through improved peptide matching. (A)** MV4-11 cells were treated with indicated concentration of FES inhibitor TAE684 for 1 h before addition of 1 or 10 µM XO44 for 30 minutes. Cells were isolated, lysed and XO44-labeled kinases were conjugated to biotin-$N_3$ using CuAAC chemistry. Proteins were precipitated using Bligh/Dyer, redissolved in PBS with 0.5% SDS and treated with avidin agarose beads to pull down labeled proteins. Beads were washed to remove unlabeled proteome and labeled proteins were eluted using Laemmli buffer containing 2 mM biotin. The proteins were separated using a 10% SDS-PAGE gel, transferred to PVDF membrane and Western blot was performed against FES (93 kDa). Arrows indicate the shift in target engagement when XO44 concentration is increased. (**B**) Including a sample with high probe dose as peptide library results in the quantification of many more kinases in the original low probe dose samples. See also Table S3.

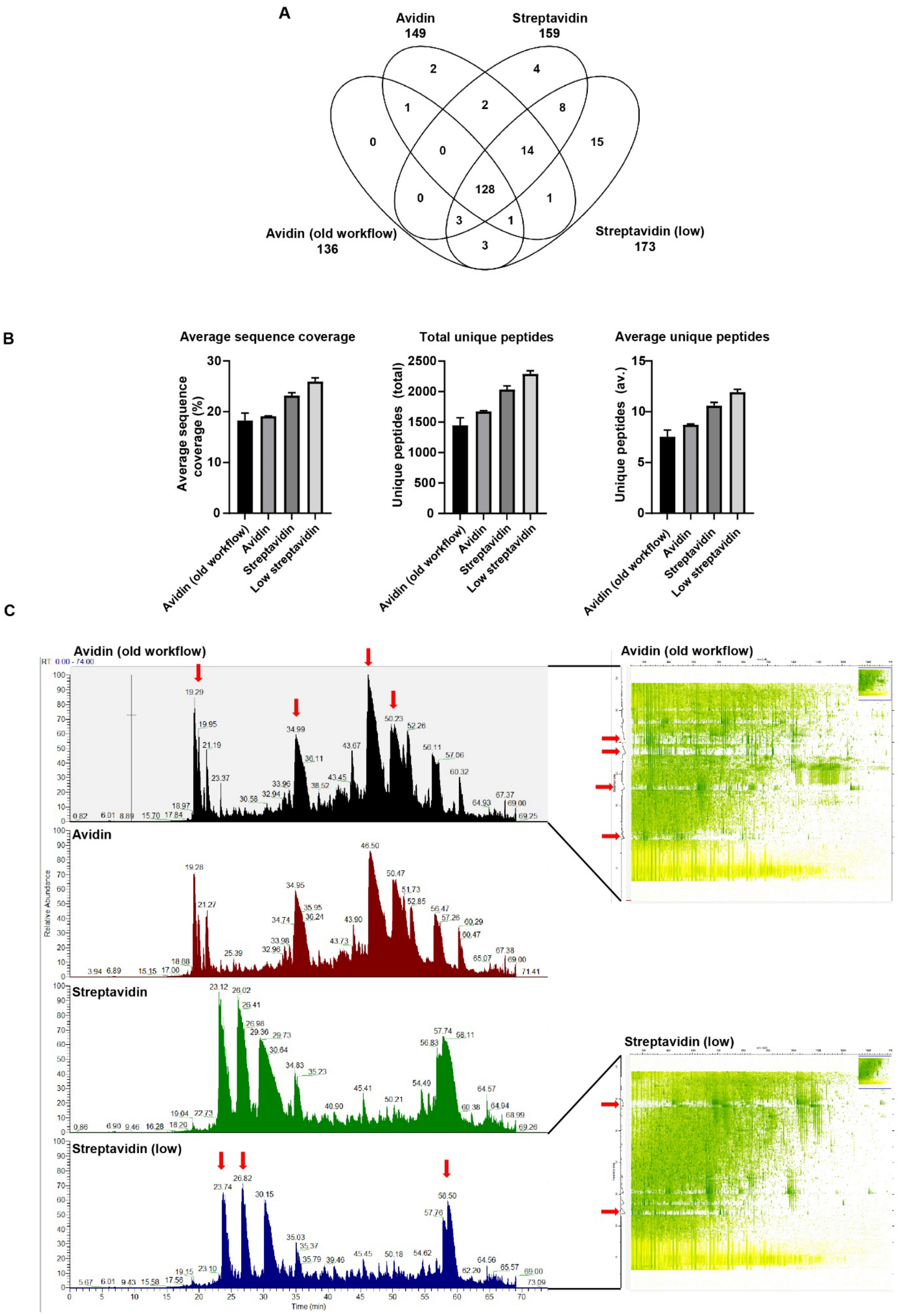
A
Avidin
149
Streptavidin
159
Avidin (old workflow)
136
Streptavidin (low)
173
2
4
1
2
8
0
0
14
15
128
0
1
3
1
3
B
Average sequence coverage
Average sequence coverage (%)
Total unique peptides
Unique peptides (total)
Average unique peptides
Unique peptides (av.)
Avidin (old workflow)
Avidin
Streptavidin
Low streptavidin
C
Avidin (old workflow)
Avidin
Streptavidin
Streptavidin (low)
Relative Abundance
Time (min)
Avidin (old workflow)
Streptavidin (low)

**Figure S5. Improvements in the chemical proteomics workflow results in lower streptavidin background, higher sequence coverage, more unique peptides and overall more kinases identified.** Pellets of XO44-treated THP-1 cells were processed for chemical proteomics using the cell line screen (old) and optimized workflow. Avidin and streptavidin agarose beads were compared in the new workflow, as well as lowering the amount of beads used. The obtained data was analyzed using MaxQuant and probe-enriched kinases were compared, defined as probe-enriched proteins annotated in Uniprot with KW-0418 of which 2+ unique peptides were found with a LFQ ratio of >2 between DMSO and XO44-treated replicates. (**A**) Overall comparison of kinases identified shows high overlap between the different workflows as well as a large gain in identified kinases. Figure generated using Venny 2.1. (**B**) With the improved workflow, the identified probe-enriched kinases show higher average sequence coverage, as well as more average and total unique peptides. (**C**) Normalized 2D and 3D chromatograms generated from Xcalibur and MaxQuant respectively show less (strept)avidin peptide contamination as well as less ion suppression, indicated by red arrows. See also Table S4.

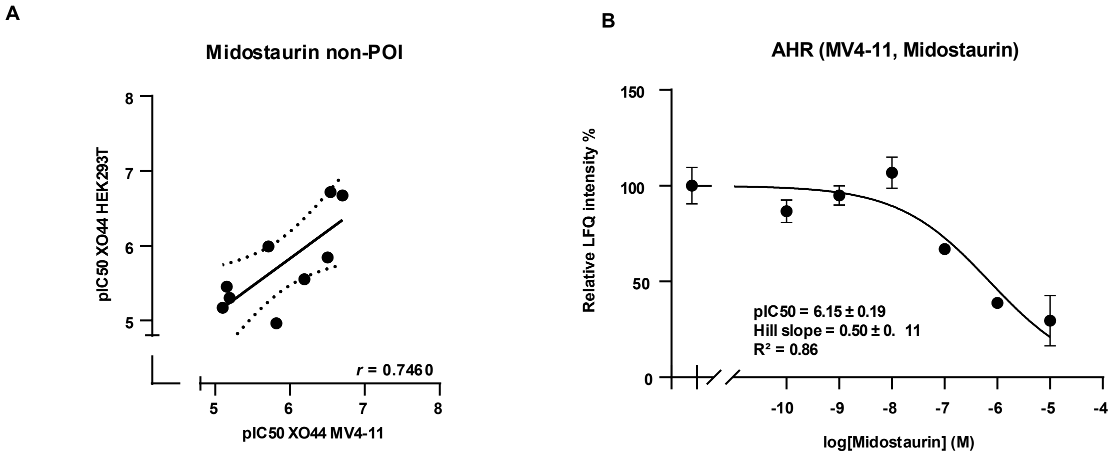


**Figure S6. Engaged proteins outside of the kinome.** (**A**) Correlation of non-kinase $pIC_{50}$ values of midostaurin between HEK293T and MV4-11 as determined by CellEKT. (**B**) Full dose response curve of AHR inhibited by midostaurin in MV4-11. Dose response curve is measured at 6 different concentrations with each concentration measured in biological duplicates n = 2. See also Table S5 and S6.
